# The Central Coupler of ClpXP Controls Intersubunit Communication and Couples Intrasubunit Chemical Energy with Force Generation

**DOI:** 10.64898/2026.04.10.717754

**Authors:** Robert P. Sosa, Alfredo J. Florez Ariza, JeongHoon Kim, Alexander Tong, Zhen-Hui Kang, Alicia Li, John Kuriyan, Carlos Bustamante

**Author notes:** These authors contributed equally: Robert P. Sosa, Alfredo J. Florez Ariza. Corresponding author: Carlos Bustamante.

## Abstract

ClpX is a clockwise hexameric helical arrangement that hydrolyzes ATP to unfold proteins and translocate them into the proteolytic chamber. We investigate the central coupler, a three α-helix module conserved among AAA+ ATPases, which is proposed to enable intersubunit communication and mechanochemical coupling. Although fundamental in AAA+ ATPases, the molecular mechanism underlying these processes remains elusive in ClpX. By combining single-molecule optical tweezers, biochemical assays and single particle cryo-EM we demonstrate that the central coupler, of the second-highest subunit of the ClpX clockwise helical arrangement, positions its Arginine-finger, sensor-I and sensor-II to contact the ATP of the anticlockwise neighbor subunit to trigger its hydrolysis. Then, the central coupler of the highest subunit rapidly couples the ATP hydrolysis with the downward motion of its substrate-translocating loop, enabling fast force generation to swiftly unfold protein substrates while maximizing the thermodynamic efficiency of the motor.

## INTRODUCTION

AAA+ enzymes (ATPases Associated with various cellular Activities) are protein complexes that convert chemical energy into mechanical work, enabling them to perform a wide range of biological functions. These functions include protein unfolding and translocation, performed by ClpX, the 26S proteasome, and YME1, or remodeling and disassembly of macromolecular complexes performed by Spastin, NSF, and Vps4^1–3^. The well-studied ClpXP motor, found in most bacteria and in mitochondria, plays crucial cellular roles, including quality control of protein folding during ribosomal translation, regulation of cellular levels of DNA repair proteins in stress response, and biosynthesis of heme precursor in mitochondria^4^. ClpXP serves as a model for understanding other AAA+ motors due to its extensive biochemical, structural, and single-molecule characterization.

ClpXP is a multi-subunit protein complex composed of a homo-hexameric ring-shaped ATPase, ClpX, and two heptameric rings of peptidase arranged in a tail-to-tail configuration to form a degradation chamber, ClpP^4^. ClpX utilizes ATP hydrolysis to recognize, unfold, translocate, and deliver the protein substrate to ClpP. Binding to protein substrates occurs by the recognition of a specific, exposed and unstructured sequences, such as the highly conserved ssrA-tag attached by the cell to proteins targeted for degradation (Fig. 1a). ClpX monomers are arranged in a helical shape, and each of them folds independently into small and large domains separated by a cleft. This cleft is lined with the Walker-B and Arginine-finger (Fig. 1b), which are involved in ATP hydrolysis and sensing of the neighboring nucleotide, respectively^4–7^. These motifs, also found in other AAA+ ATPases, including the Walker-B^8–10^ and others like the IGF-loops involved in assembly with ClpP^10,11^, the RKH-loops involved in substrate recognition^12,13^, and the pore loops involved in polypeptide translocation^10,12,14,15^, have been identified in ClpX and investigated extensively through biochemical and single-molecule techniques^9,14,16–19^.

**Fig. 1.**
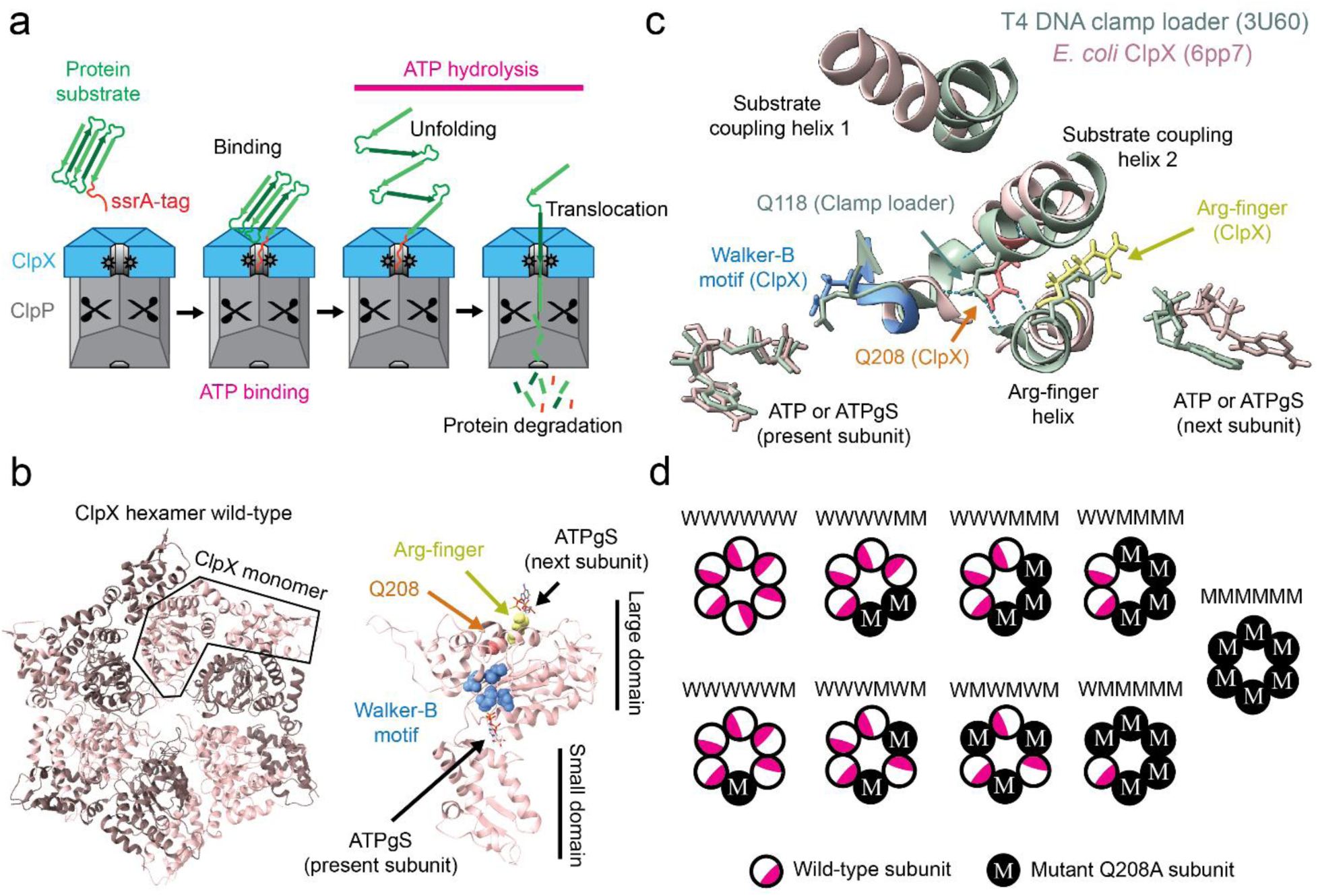
Protein unfolding by ClpXP, architecture of central coupler. **a)** Cartoon of the general mechanism of action of ClpXP. **b)** ClpX hexamer (left), showing the general context of Glu208 within a ClpX subunit (right). The Arg-finger, Q208 and walker-B of ClpX are shown in yellow, orange and blue, respectively. **c)** Comparing the structural alignment of the α-helices of the central coupler, the Walker-B, and the two ATP(γS) in contact with those elements of ClpX (6PP7 PDB, pink) with respect to those same elements in the T4 DNA clamp loader (3U60 PDB, calm green). The Arg-finger, Q208 and Walker-B of ClpX are shown in the same color code as in b. The Q118 of the T4 DNA clamp loader is indicated. H-bonds formed by Q118 and Q208 are shown as dotted lines. **d)** Cartoon of the representative ClpX hexamers with wild-type (W) subunits and harboring the mutation Q208A (M).

Subramanian *et al*.,^20^ recently conducted a high throughput deep mutagenesis screen to map the mutational sensitivity of each amino acid in another member of the AAA+ ATPase family, the T4 DNA clamp loader. This study revealed that this protein is highly sensitive to mutagenesis in Q118 (Q208 in ClpX). Furthermore, structural analysis and simulations revealed that Q118 is not involved in catalysis but instead forms a H-bonded junction in an alpha helical-containing unit that the authors named the “central coupler”. Q118 maintains the rigidity of the central coupler, which connects the catalytic centers of the clamp loader to its DNA substrate (Fig. 1c). Interestingly, this structural feature is also present in ClpX and in other AAA+ ATPases^20^. The authors suggested that, despite their extraordinary functional diversity, the rigidity of the central coupler may be the architectural feature responsible for cooperative ATP-driven action among neighboring subunits in AAA+ ATPases^20^.

Here, we study the critical role of the rigidity of the central coupler of ClpXP, combining biochemical, high-resolution optical tweezers, and single-particle Cryo-EM methods. Our results shed light on the molecular mechanism of intersubunit coordination and mechanochemical coupling during force generation by ClpXP.

## RESULTS

### The central coupler of the T4 DNA clamp loader and ClpX

The central coupler of the T4 DNA clamp loader is composed of three alpha helices: (i) the Arg-finger helix, which contains the residue that senses the ATP status of the neighboring anti-clockwise subunit; (ii) the substrate-coupling helix 2, where Q118 is located, which establishes the H-bonded network linking the catalytic Walker-B motif and the Arg-finger helix; and (iii) the substrate-coupling helix 1 (Fig. 1c), which is connected to the DNA substrate^20^. Alignment of the central couplers, the Walker-Bs, the ATP(γS) nucleotides of a subunit and that of its clockwise neighbor of the T4 DNA clamp loader with the same elements of *E. coli* ClpX show that they share a similar architecture (RMSD = 1.1 Å). Importantly, the relevant glutamine Q208 of ClpX is structurally similar to Q118 of the clamp loader (Fig. 1c). However, the substrate-coupling helices 1 of both structures are the most divergent. This result is expected because these helices are connected to their different respective subtrates, dsDNA for the clamp loader and an unfolded polypeptide for ClpX.

Simulations of the T4 DNA clamp loader^20^ indicate that mutation Q118A eliminates H-bonds that link the walker B motif and the Arg-finger helix. Accordingly, we introduced the corresponding mutation (Q208A) in the ClpX monomer. Among other possible residues forming H-bonds in the central coupler of ClpX, Q208A disrupts the network of H-bonds (Fig. 1c), weakening its interaction with the Walker-B and Arg-finger helix, resulting in a flexible central coupler.

### ClpX monomer mutant fails to assemble into active hexamers

The mild mutation of the glutamine residue (Q118N) in the T4 DNA clamp loader suggest that it may disrupt hexamer assembly^20^. We tested whether Q208A ClpX monomers (mono^QA^) could self-assemble into active hexamers. Wild-type monomers (mono^WT^) self-assemble, resulting in a broad peak in the size exclusion chromatogram, with the earliest region corresponding to the hexamer form and later regions corresponding to lower-order oligomerization states (Supplementary Fig. 1). In contrast, the chromatogram of mono^QA^ showed no absorbance around the hexamer peak, indicating that mutant monomers are unable to form hexamers (Supplementary Fig. 1). However, single-chain constructs (MMMMMM and WWWWWW, see below) displayed a similar peak as the mono^WT^ hexamer (Supplementary Fig. 1). Consistently, a degradation assay of Titin^CM^-ssrA using SDS-PAGE revealed that hexamers formed by mono^WT^ and WWWWWW were active, while MMMMMM was active but significantly slower, and mono^QA^ showed no activity (Supplementary Fig. 1, inset). Therefore, the disruption of the H-bonded junction by Q208A in ClpX monomers results in their inability to self-assemble into active hexamers. To distinguish defects in oligomerization from ClpX functional activities, we studied the single-chain ClpX mutant constructs (Fig. 1d).

### Single-chain ClpX constructs

We introduced the Q208A mutation in the covalently-linked subunits of single-chain ClpX hexamers^21^. This latter architecture allows us to (i) precisely control the number and spatial positioning of Q208A subunits (Fig. 1d) and (ii) study processes related to the mechanochemical cycle (ATP hydrolysis, coupling and force generation) and downstream functional activities (polypeptide translocation and protein unfolding) of fully assembled ClpX hexamer without oligomerization defects. In these constructs we refer to subunits as either “W” (for wild-type) and “M” (for subunit harboring the mutation Q208A). For example, WWWWWM and WWWMMM refer to constructs with a single-isolated mutant or occupying the three contiguous positions in the single-chain hexamer, respectively. WWWWWW, WWWWWM, WWWMWM and WMWMWM reveal the effect of adjacent wild-type and mutant subunits relative to two adjacent mutant subunits in WWWWMM, WWWMMM, WWMMMM (Fig. 1d). Comparing the activity of WWWMMM vs WMWMWM provides the most contrasting effect of the configuration of mutant subunits.

### Single-molecule optical tweezers set-up

To investigate the implications of a flexible central coupler by the Q208A mutation, we used dual-trap high-resolution optical tweezers^14,16,17^ to follow the activity of single-chain ClpX hexamers in complex with ClpP (ClpXP) during force generation, polypeptide translocation, and protein unfolding^9,14,16–19^. The protein substrate has a biotin in the N-terminus, followed by a GFP moiety, four unstructured titin-I27 (4xTitin^CM^) and ends in a ssrA-tag at the C-terminus (Fig. 2a). We immobilized C-terminally biotinylated ClpX hexamers onto the surface of 2 µm polystyrene beads via biotin-streptavidin linkages. The substrate was attached to another 2 µm polystyrene bead using a neutravidin-biotin-DNA handle bridge (Fig. 2a). Both beads were held in separate optical traps. After bringing the two beads close together, ClpXP engages the ssrA-tag, forming a tether between them (Fig. 2a). The molecular trajectories obtained with WWWWWW reproduced those previously observed and reported^9,14,16–19^. Starting the assay at a tension of 6 pN in the presence of ATP, while holding the trap positions fixed (*passive mode*), ClpXP begins to translocate the 4xTitin^CM^ moieties which results in a corresponding increase in tension opposing the motor. Translocation was monitored by a gradual decrease in the distance between beads held in the fixed traps (Fig. 2b). Upon encountering GFP (the polypeptide chain is threaded from the top of the ClpX pore to the proteolytic chamber of ClpP), ClpXP pauses as it attempts to unfold the protein, which then results in a sudden increase of bead-to-bead distance once successful. This tether increment is followed by another gradual decrease of inter-bead extension reflecting ClpXP translocation over the unfolded GFP polypeptide (Fig. 2b). As reported, ClpXP translocation traces are made up of periods of dwells where the motor does not move and interrupted by translocation steps^9,14,16–19^, which we determined using the Kalafut-Visscher step finding algorithm (see methods, Supplementary Fig. 2). We validated the robustness of the algorithm with simulations (Supplementary Fig. 3, 4).

**Fig. 2.**
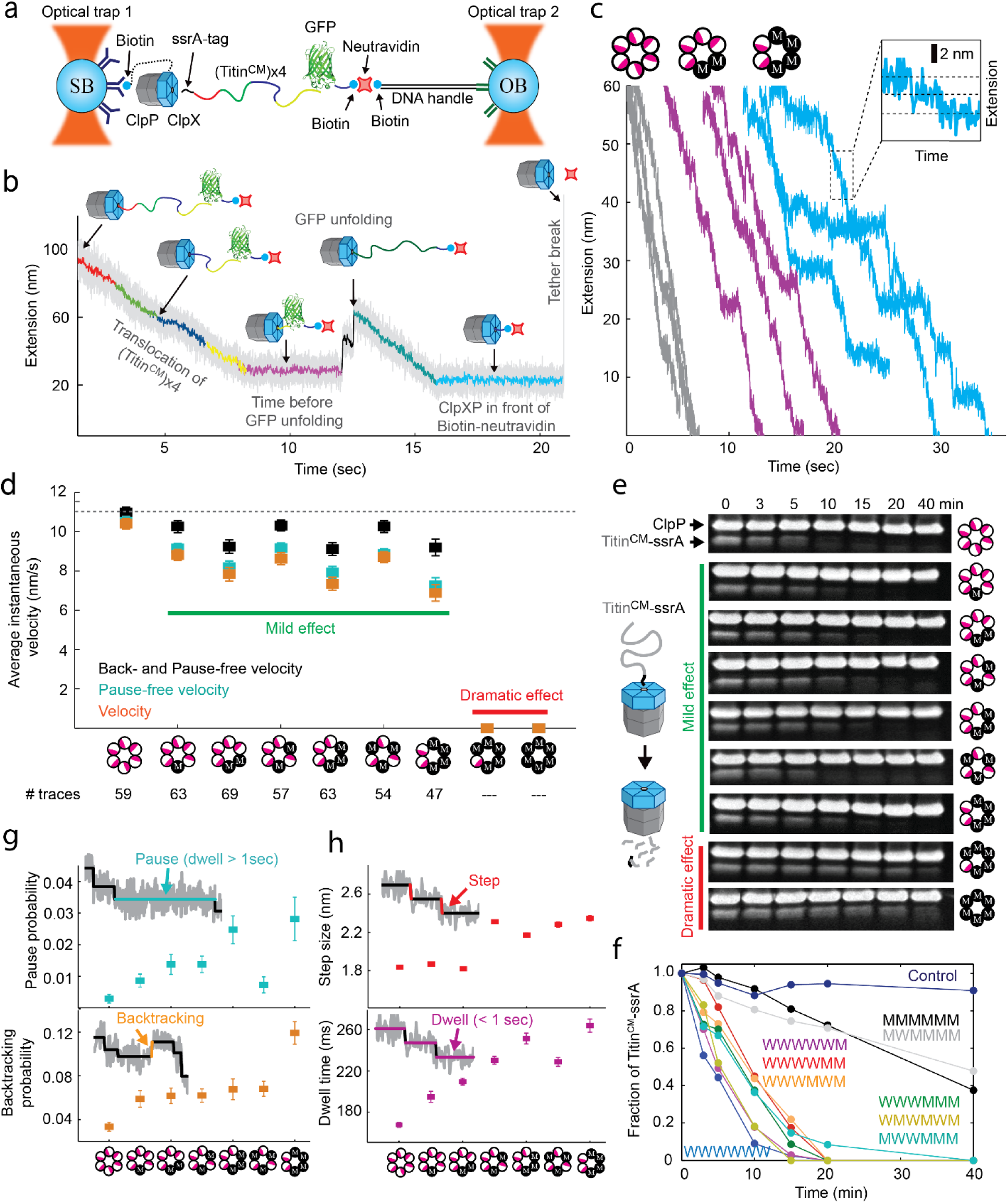
Optical tweezers set-up and effect of Q208A mutation on polypeptide translocation. **a)** Experimental set-up in the dual trap high-resolution optical tweezers. **b)** Single-molecule trajectory (extension vs time) in passive mode of polypeptide translocation and protein unfolding at 2.5 kHz (grey) and down-sampled to 100 Hz (color). **c)** Representative examples of polypeptide translocation at 100 Hz of wild-type, WWWWAA, and WWAAAA. **d)** Plot of the total velocity, pause-free velocity, pause- and backtracking-free velocity (nm/s) of polypeptide translocation of wild-type and Q208A hexamer mutants. Error bars represent the *SEM* = *SD*/√*p*, where *p* is the number of data points. WWWWWW (N = 59 traces), WWWWWM (N = 63), WWWWMM (N = 69), WWWMWM (N = 57), WWWMMM (N = 63), WMWMWM (N = 54), and WWMMMM (N = 47). **e)** SDS-PAGE gel degradation of Titin^CM^-ssrA by different Q208A mutants. **f)** Plot of fraction of substrate remaining (Densiometric analysis) vs degradation time (min) of all constructs. Control is all the components of reaction but without ClpX. **g)** Top, backtracking probability. Bottom, pause probability per trace and per Q208A hexamer mutants. Error bars = SEM. **h)** Average step size (nm) and dwell time (ms) during translocation, excluding pauses and backtracking, for WWWWWW (2829 dwells-steps), WWWWWM (2973 dwells-steps), WWWWMM (3061 dwells-steps), WWWMWM (1991 dwells-steps), WWWMMM (2107 dwells-steps), WMWMWM (1330 dwells-steps), and WWMMMM (1556 dwells-steps). Error bars = SEM.

### A flexible central coupler does not affect translocation over unstructured polypeptides

Single-molecule trajectories were subjected to step analysis to extract dwell times and step sizes (see methods). Instantaneous velocity values were determined by dividing the magnitude of the step sizes by the dwell time preceding that step. WWWWWM, WWWWMM, WWWMMM, and WWMMMM mutants displayed only a minor effect on their average instantaneous translocation velocity over unstructured polypeptides, compared to that of WWWWWW (Fig. 2c, d). A similar result is observed for other configurations of double (WWWMWM) and triple (WMWMWM) mutants (Fig. 2d). Also, WMMMMM and MMMMMM while able to engage with the ssrA-tag, as evidenced by the formation of a stable tether between the optical traps, they were unable to translocate, even when the opposing tension was lowered from ∼6 to ∼1 pN (Fig. 2d). The latter suggests that these mutants, with one or no wild-type subunits, are highly sensitive to force. The SDS-PAGE-based bulk degradation assay confirmed that WWWWWM, WWWWMM, WWWMMM, WWMMMM, and the mutants with a different configuration (WWWMWM and WMWMWM) have no impairment in their ability to degrade the Titin^CM^-ssrA substrate while WMMMMM and MMMMMM also showed a dramatic effect on degradation (Fig. 2e, f). Overall, a minimum of two subunits having a rigid central coupler, independent of their location within the hexamer, appears to be sufficient for near-optimal processive translocation of an unfolded polypeptide. Finally, the above single-molecule and bulk results also indicate that polypeptide translocation rather than substrate recognition is the rate-limiting step in the activity of ClpXP when the client protein is unfolded (Titin^CM^-ssrA), a result consistent with previous bulk studies^22^.

The molecular trajectories reveal that WWWWWW also makes infrequent pauses (defined when dwell time longer than 1 sec^14,16,17^, see Fig. 2g). Traces also displayed backward steps, which we refer to as “backtracking”, because they are of comparable size to translocation steps (Fig. 2f, Supplementary Fig. 5a). However, they are much shorter than events of large tether increase (slips) that are likely due to temporary grip loss of the client polypeptide by ClpX^16^. Although rare, the probability of both pauses and backtracking events, as well as pause duration, increase significantly from single to quadruple mutants in a subunit arrangement-independent manner (Fig. 2g). Thus, the rigidity of the central coupler also appears to play a role in the polypeptide translocation processivity of ClpXP.

### The central coupler enables intersubunit communication and mechanochemical coupling

As reported previously^14^, during polypeptide translocation, ADP release and ATP binding take place during the dwell phase whereas ATP hydrolysis and phosphate release coincide with the stepping/burst phase^14^. As expected from the near-optimal translocation velocity observed with up to four mutant subunits over a the fully unfolded Titin^CM^-ssrA substrate, (Fig. 2d), we find that the average step size of the motor is also invariant to the incorporation of these mutations, remaining nearly fixed at ∼2 nm (Fig. 2h, Supplementary Fig. 5a), with the differences in velocity coming from lengthening of the dwells (Fig. 2h, Supplementary Table 1). The pair-wise distribution analysis (see methods) confirms that ClpX performs translocation steps of ∼2 nm (Supplementary Fig. 6).

ClpXP translocates ∼1 nm per ATP hydrolyzed^14^; we thus wondered whether Q208A affects the ATP hydrolysis rate of the enzyme and, therefore, the coupling efficiency between chemical energy consumed and the motor’s mechanical output (i.e., its mechanochemical coupling). To address this question, we used an ATPase assay to determine the maximum ATP hydrolysis rate (k_cat_), the Michaelis-Menten constant (K_m_), and the Hill coefficient (n_H_) of hexamers. To ensure continuous polypeptide translocation, the fully unfolded substrate Titin^CM^-ssrA was supplied at a saturating concentration of 500 μM compared to 0.4 μM of ClpX.

The ATPase assay of all hexamer mutants tested in the optical tweezers (Fig. 3, Supplementary Fig. 7a) revealed that WWWWWW and WWWWWM have similar values of k_cat_, that WWWWMM, WWWMWM, WWWMMM, and WWMMMM have ∼1.5-fold larger k_cat_ values than WWWWWW, and that the triple variant WMWMWM has the highest k_cat_ value, ∼4-fold higher (Fig. 3a). The K_m_ followed a similar trend (Fig. 3b). A linear relationship between K_m_ and k_cat_ is to be expected if Q208A does not change the nucleotide binding constant (K_D_) but increases k_cat_, according to the equation K_m_ = K_D_ + (1/k_on_)k_cat_. Indeed, in most cases K_m_ and k_cat_ are linearly related, confirming that the loss of the H-bonded junction by the incorporation of Q208A raises the Km by increasing the corresponding k_cat_ of ATP hydrolysis, without affecting ATP binding (Fig. 3c). The only outlier in Fig. 3c is the pentamutant, whose Km and kcat cannot be estimated reliably since it is not possible to reach saturation in the ATPase assay (Supplementary Fig. 7b). The Km was much higher in the case of MMMMMM (Supplementary Fig. 7b). To have a further confirmation that K_D_ is also invariant in this case, we measured ATP binding via a fluorescence enhancement assay^23^ (see methods) and found that WWWWWW and MMMMMM have identical ATP binding curves (Supplementary Fig. 7b inset). Thus, Q208A increases the K_m_, via its effect on the k_cat_ without modifying the dissociation constant of the nucleotide. Taken together, these results indicate that Q208 in ClpX down regulates the ATP hydrolysis rate of the hexamer.

**Fig. 3.**
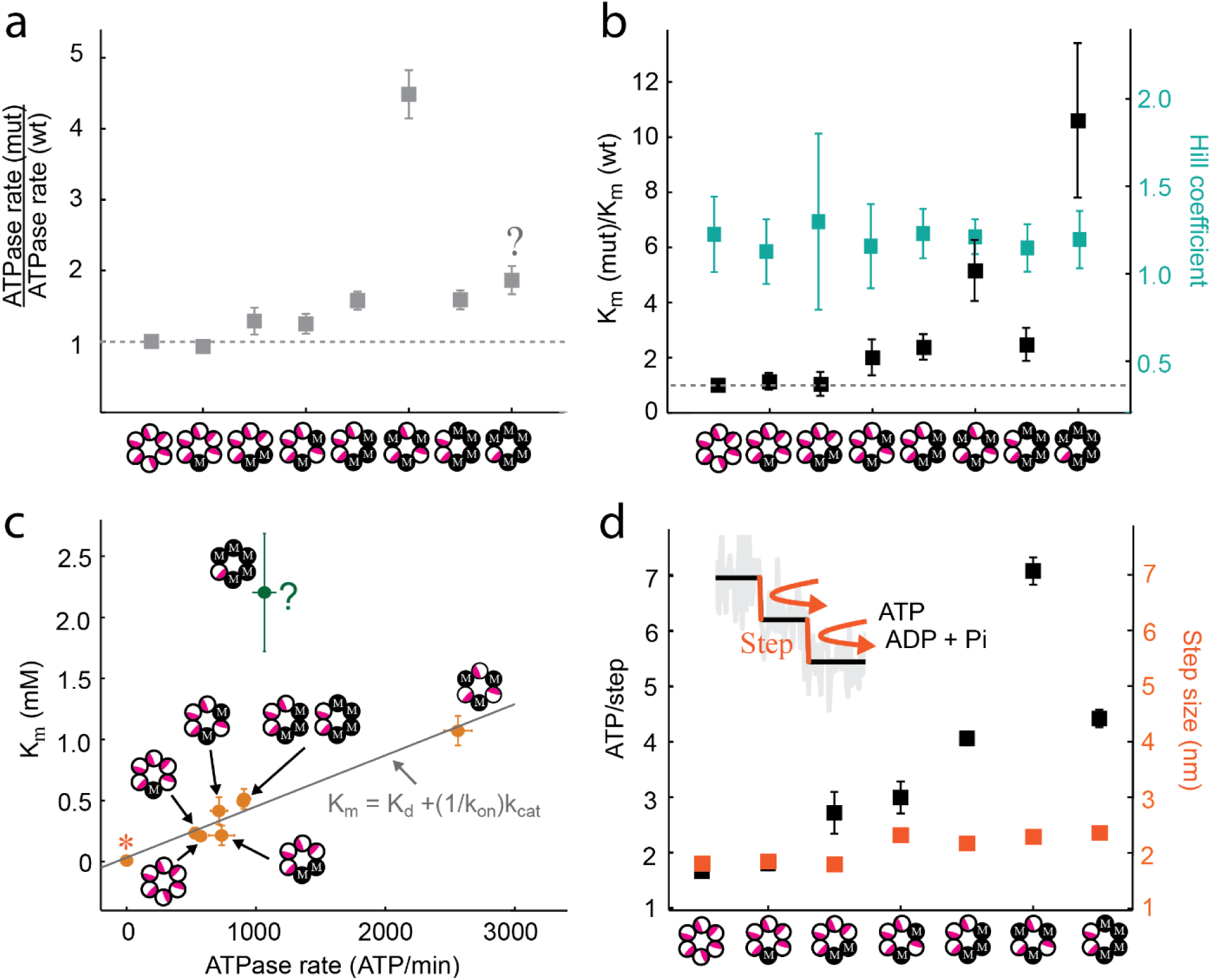
Effect of the Q208A mutation in the mechanochemical coupling of ClpXP. **a)** Ratio of Maximum ATP hydrolysis rate (kcat, ATP/min) of Q208A hexamer mutants relative to of the wild-type. **b)** Ratio of Km values of Q208A hexamer mutants relative to of the wild-type. Each ATP condition per construct was carried out via triplicate (n = 3). Hill coefficient of wild-type and Q208A hexamer mutants. **c)** Plot of Km values (mM) vs kcat (ATP/min) wild-type and Q208A hexamer mutants. **d)** Comparison of number of ATP consumed per translocation step (ATP/step) and the average step size of wild-type and Q208A hexamer mutants.

The efficiency of the motor is equivalent to the number of ATP molecules consumed per step. It can be obtained by dividing k_cat_ (ATP/s) by the number of steps per second. The consumption of ATP per step mostly increases monotonically with the number of mutants in the hexamer. WWWWWW and WWWWWM consume ∼2 ATPs/step, WWWMWM and WWWWMM spend ∼3 ATPs/step, whereas WWWMMM and WWMMMM hydrolyze ∼4 ATPs/step. Notably, WMWMWM consumes ∼7 ATPs/step (Fig. 3d). Also, WMWMWM hydrolyzes ATP ∼2-fold times faster than WWMMMM and WWWMMM, Fig. 3d) indicating that not only the number of mutants but also their spatial arrangement (intersubunit communication) in the hexamer determines the efficiency of the motor. In summary, weakening the rigidity of the central coupler reduces the mechanochemical coupling of ClpX, i.e., its transduction of chemical energy into mechanical work, resulting in a larger number of hydrolyzed ATPs to accomplish the same mechanical task. Additionally, our results also suggest that efficient mechanochemical coupling requires appropriate intersubunit communication (Supplementary table 5).

### The rigidity of the central coupler is essential to overcome high mechanical barriers

Protein unfolding is the rate-limiting step during proteolysis by ClpXP because folded proteins pose a greater mechanical barrier for the motor than translocation over an unfolded polypeptide^24,25^. To establish the role of the central coupler’s rigidity on the unfolding process, we investigated the ability of the various Q208A hexamers to unfold GFP (Supplementary Fig. 8). The probability of unfolding was calculated as the fraction of single-molecule traces displaying GFP unfolding events^14,16,17^ (Fig. 4a). The single mutant had similar unfolding probability to wild-type, but WWWWMM and WWWMWM lowered the unfolding probability from ∼75% to ∼45%, whereas WWMMMM and WWWMMM displayed a drastic decrease of unfolding probabilities down to ∼25% (Fig. 4b). Notably, WMWMWM showed 0% unfolding probability (Fig. 4b, c).

**Fig. 4.**
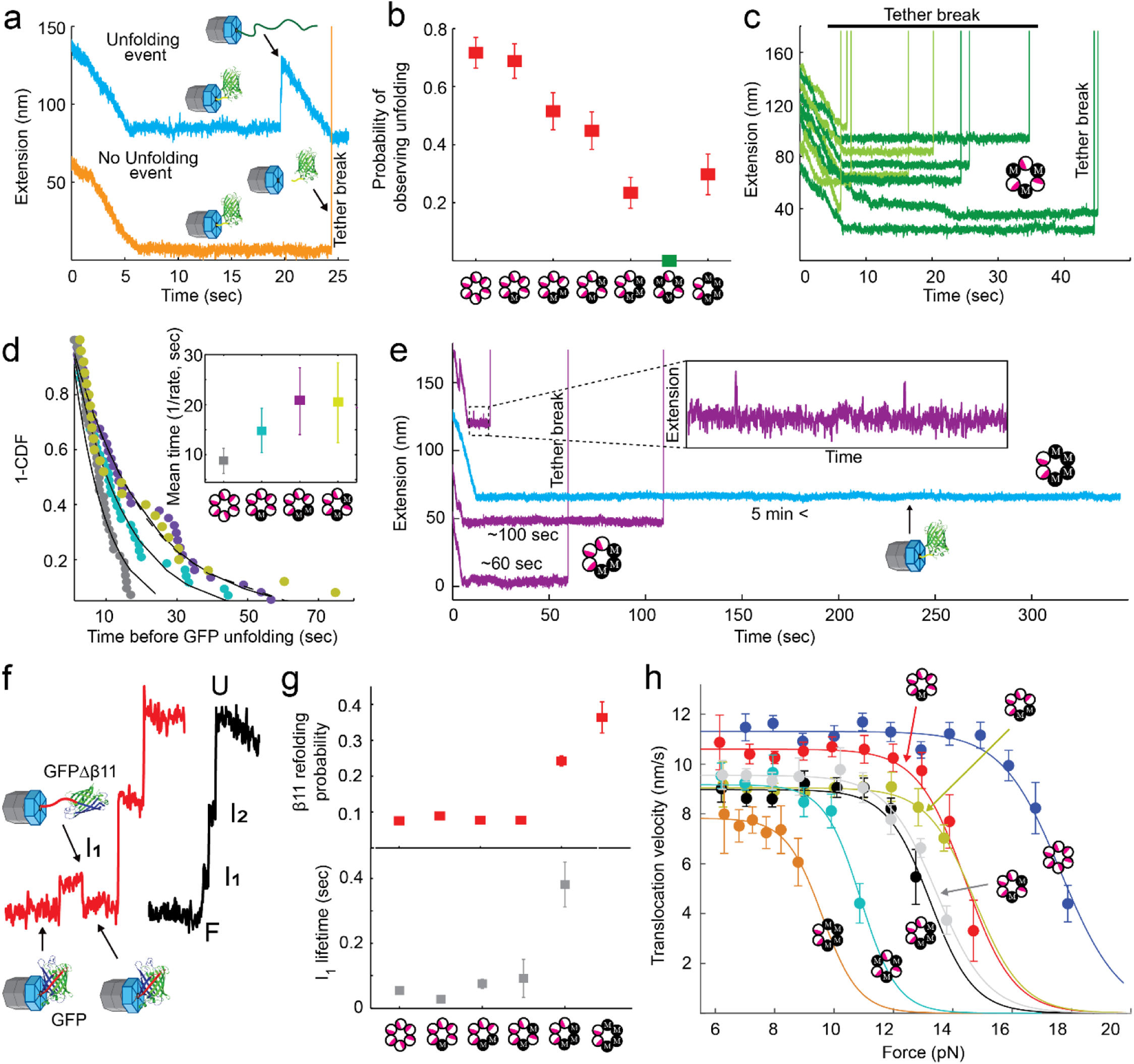
Effect if the Q208A mutation in the capacity of ClpX to overcome high-mechanical barriers. **a)** Representative traces of ClpX activity with and without unfolding event. **b)** Probability of observing unfolding events out of total traces per Q208A hexamer mutants. WWWWWW (n = 42 GFP unfolding events out of N = 59 traces), WWWWWM (n = 43 of N = 63), WWWWMM (n = 37 of N = 69), WWWMWM (n = 26 of N = 57), WWWMMM (n = 14 of N = 63), WMWMWM (n = 0 of N = 54), and WWAAAA (n = 13 of N = 47). Error bars are calculated with the boostrapping method. **c)** Representative examples of single-molecule traces of the WMWMWM mutant. **d)** Inverse of the cumulative density function plot of the time to unfold GFP by WWWWWW, WWWWWM, WWWWMM and WWWMWM. Dots are experimental data, and continuous line represent the fitting to a single exponential equation to extract the kinetic lifetime. Inset: mean time of GFP unfolding of WWWWWW, WWWWWM, WWWWMM and WWWMWM. Error bars = SEM. **e)** Representative examples of single-molecule traces of WWWMMM and WWMMMM mutants. **f)** Representative examples of the failed and successful attempt to overcome the first intermediate during GFP unfolding by ClpX. **g)** Top, plot of the β11 refolding probability. Bottom, plot of the average I1 lifetime of WWWWWW (I1 is present is 25 unfolding events), WWWWWM (16 events), WWWWMM (13 events), and WWWMMM (5 events). Error bars = SEM. **h)** Translocation velocity (nm/s) vs force (pN) of the Q208A hexamer mutants. Error bars = SEM.

When ClpXP arrives at the GFP, the protease spends some time attempting to mechanically unfold the protein (Fig. 4a). We found that the pause before GFP unfolding increased from ∼9 sec for WWWWWW to ∼15 sec for WWWWWM, and to ∼21 sec for WWWWMM (Fig. 4d). Notably, a double mutant with different hexamer configuration (WWWMWM) also took ∼21 sec to unfold GFP. This trend suggests that WWWMMM and WWMMMM would require even a larger time to unfold GFP (>> 21 sec). However, it was not possible to quantify the unfolding time in these mutants, since they took such long time to unfold the GFP (> 5 min) that the tether often broke before the unfolding event was observed (Fig. 4e). Overall, optimal mechanical operation and force generation require all six subunits to have wild-type central couplers. Consistent with this conclusion, we found that a single Q208A mutation dramatically decreases the ability of wild-type ClpXP to pull the biotin out of its binding pocket in neutravidin (Supplementary Fig. 9).

We previously determined^17^ that the initial extraction of the C-terminal β-strand 11 (β11) of GFP represents the first mechanical barrier for successful unfolding. The mean refolding time constant of β11 into GFP was determined to be ∼230 msec at low (< 200 µM) ATP concentrations^17^. Assuming this is true in our case, because we used similar experimental conditions with saturating ATP, if ClpX fails to translocate over the unfolded β11 in less than 230 msec, this element is likely to refold, and ClpX will have to start a new unfolding attempt. The extraction of β11 is visible as an increase in extension to a state called I1 (Fig. 4f, Supplementary Fig. 8). The dwell time during translocation is ∼165 msec for WWWWWW and below 225 msec for WWWWWM, WWWWMM and WWWMWM, which is less than the β11 refolding time (Fig. 2g), and hence only 10% of the time β11 is seen to refold. In contrast, the dwell time of WWWMMM and WWMMMM are ∼250 msec and ∼270 msec, respectively (Fig. 2g), longer than the β11 refolding time, hence these mutants display a refolding event ∼25% and ∼35% of the time, respectively (Fig. 4g). Furthermore, for the case in which extraction of β11 leads to the full unfolding of GFP, the mean average time of this intermediate remains below 100 msec in WWWWWW, WWWWWM, and WWWWWM. In contrast, this value increases to ∼400 msec in WWWMMM (Fig. 4g). Altogether, these results indicate a rigid central coupler favors the quick conversion of chemical energy from ATP hydrolysis into mechanical displacement on the unfolded substrate allowing ClpX to effectively invade it before it refolds.

### A rigid central coupler determines the maximum generation of force

The results above suggest that wild-type central coupler across all subunits are required for ClpXP to generate force efficiently and rapidly to overcome high mechanical barriers. To have direct evidence of this hypothesis, force-velocity curves were obtained for the wild-type and various configurations of the Q208A mutation in the hexamer (Fig. 4h). These experiments are performed in passive mode in which the traps positions are held fixed. As the motor translocates over the unfolded polypeptide, the tension in the tether increases. Eventually, the translocation velocity drops to zero when the external load equals the maximum force the motor can generate (its stall force)^14,16^. The force dependence of the pause-free velocity can be fit to a single-barrier Boltzmann equation (see methods), from which it is possible to obtain the external force F_1/2_ that reduces the translocation velocity of the motor to one-half of its maximum value^26^.

WWWWWW and WWWWWM showed comparable velocities at zero force, but their F_1/2_ were ∼19 and ∼14 pN, respectively (Fig. 4h). Interestingly, WWWWMM exhibited a similar F_1/2_ ∼14 pN as WWWWWM, but with slightly smaller zero-force velocity (∼9 nm/s vs. ∼11 nm/s, respectively). WWWMMM increased motor’s sensibility to force; F_1/2_ ∼13 pN, while WWMMMM has a drastic reduction in F_1/2_ down to ∼ 9 pN. Furthermore, WWWMWM is more sensitive to force than WWWWMM, showing an F_1/2_ of 13 pN vs ∼14.5 pN, whereas WMWMWM and WWWMMM showed F_1/2_ of ∼13.5 pN and ∼10.3 pN, respectively (Fig. 4h). Thus, while the translocation velocity of WWWWMM, WWWMWM, WWWMMM, WMWMWM and WWMMMM extrapolated at zero force over an unfolded polypeptide did not deviate significantly from ∼9 nm/s, F1/2 ranged between ∼15 pN and ∼9 pN (Fig. 4h). Altogether, these results provide direct evidence that the rigidity of the central coupler is required for ClpX to generate maximum force (∼21 pN) to overcome high mechanical barriers (protein unfolding and complex disassembly). Conversely, the rigidity of the central coupler appears not to be essential for ClpX to overcome low mechanical barriers (translocation of an unfolded polypeptide).

Finally, these results also explain why WMMMMM and MMMMMM are not active in the optical tweezers. First, the more subunits harboring the Q208A mutation, the more sensitive is the ClpX hexamer to external forces. Second, although WWMMMM have a mild reduction in polypeptide translocation (in SDS-PAGE assay and by its activity below 8 pN in tweezers), F1/2 is reduced to ∼9 pN. Since WMMMMM and MMMMMM display a dramatic effect on Titin^CM^-ssrA degradation in the SDS-page assay (Fig. 2d), it is not surprising that those ClpX hexamers will be highly sensitive to external force in the optical tweezers.

### Cryo-EM structure of WMWMWM ClpXP complexes

To establish the structural basis behind the crucial role of the central coupler, we used single particle cryo-EM (Supplementary Fig. 10a) to study the WMWMWM in complex with ClpP and a ssrA-tagged GFP (a roadblock for this mutant) in the presence of an ATP regeneration system (ATP-RS or a combination of ATP-RS/ATPγS). WMWMWM was chosen because it has a similar translocation velocity than the WWWWWW (Fig. 2d) but showed dramatic effects on mechanochemical coupling (Fig. 3d), and protein unfolding along with force generation (Fig. 4h).

After performing single-particle analysis, a total of three major ClpXP structural classes were obtained (Fig. 5a-c, Supplementary Data 1, Supplementary Table 2, 3). As reported in previous cryo-EM studies^6,7,27–31^, the ClpX subunits in the three classes adopt a clockwise right-handed helical or spiral conformation (A^W^B^M^C^W^D^M^E^W^F^M^ from top to bottom, with the ‘seam’ between subunit F and A) that sits on top of one of the two ClpP homo-heptameric rings, contacting it via the IGF loops (Fig. 5a-c). Class-I structure was solved at 3.7 Å nominal resolution and represents a substrate-free/nucleotide-bound ClpXP complex (Fig. 5a, Supplementary Fig. 10b-e, 11). Class-II and Class-III were obtained at 3.7 Å and 3.4 Å nominal resolution, respectively, and both represent substrate-bound/nucleotide-bound ClpXP complexes (Supplementary Fig. 10b-e, 11). Class-II was obtained in the presence of ATP-RS/ATPγS system, whereas class-III, in the presence of ATP-RS only (see methods). In all classes, the attained local resolution allowed us to identify the specific subunits harboring the wild-type Q208 (W) or the mutated A208 (M) residue as well as the state of the bound nucleotide, ATP or ADP (Supplementary Fig. 10c, d). At different threshold levels, Class-II and Class-III display an extra density on top of ClpX (Supplementary Fig. 10b), indicating that the motor is stalled in front of a mechanical barrier of a folded or partially folded structure. Although we cannot determine the sequence of aminoacids of the polypeptide within the ClpX pore (modeled as poly-Alanine), it is clearly threaded completely to the bottom subunit of the spiral. This feature is also found in other cryo-EM structures where *E. coli*’s ClpXP is reported to be stalled, presumably applying force, in front of a folded structure that can or cannot be modeled^7,28^ (Supplementary Fig. 12).

**Fig. 5.**
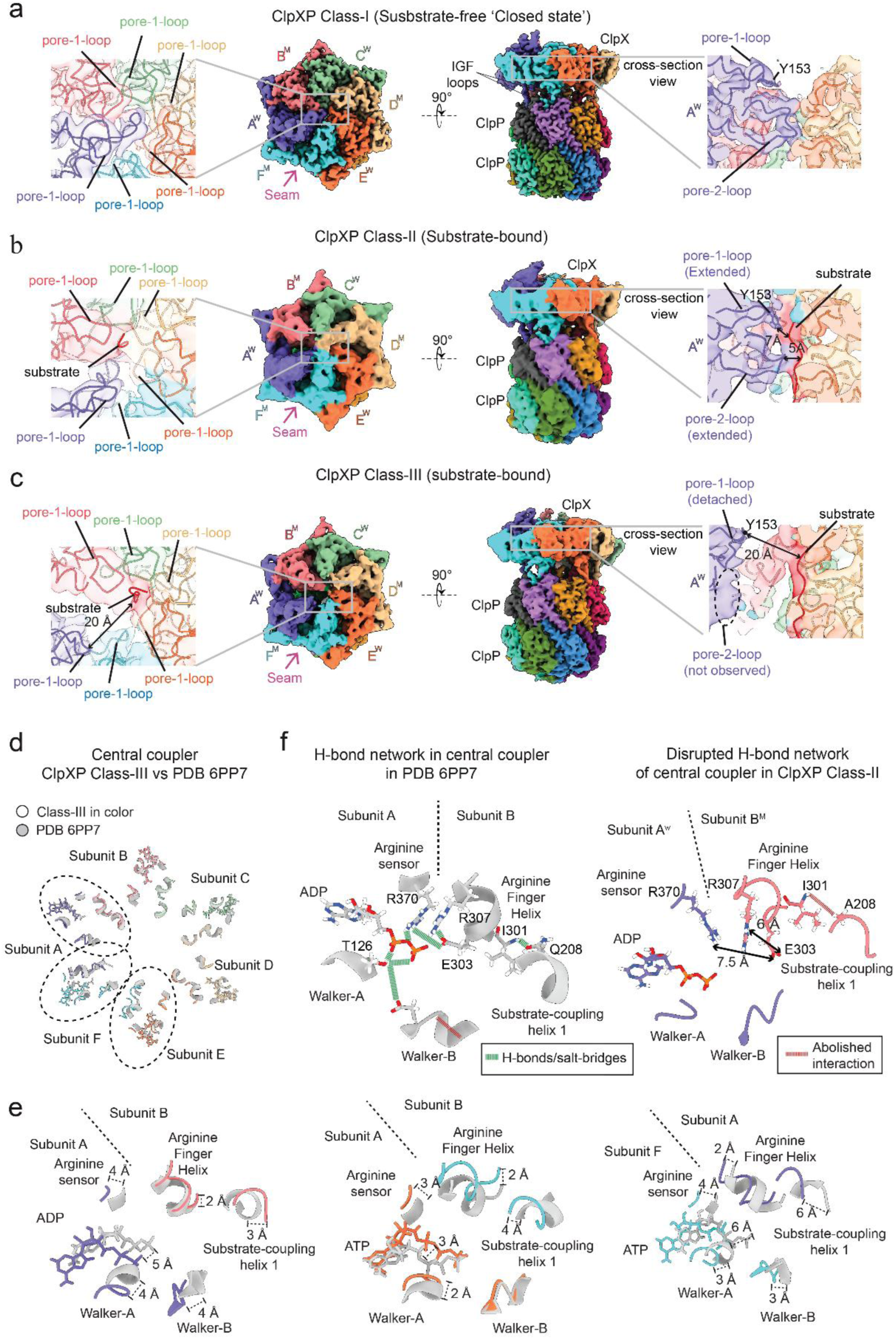
ClpXP Cryo-EM structures of the WAWAWA mutant. **a)** Top and side views of the Cryo-EM density maps of the Class-I, -II and -III ClpXP complexes. Each ClpX subunit harboring the wild-type (W) Q208 or mutant (A) A208 residue, is indicated by the corresponding superscript. See Supplementary Fig. 10d for details. **b)** Internal structural details of the top view for each ClpXP class in (A). The ClpX subunits are assembled in clockwise sense from A (top) to F (bottom) (color key indicated in figure), following the spiral arrangement of the ClpX pore-1 loops (indicated). The “seam” of the spiral (purple arrow) is shown in a. **c)** Internal structural details of the ClpX central pore region for ClpXP Class-I to -III. The Class-I is observed in a substrate-free state, while the Class-II and -III are both substrate-bound states. **d)** Visualization of the Central-Coupler elements of all the ClpX subunit interfaces after overlapping the ClpXP Class-III vs PDB 6PP7. The subunit interfaces showing significant rearrangements are highlighted with dashed ovals. ClpX subunits of Class-III are colored as in (A), while ClpX PDB 6PP7 is colored in gray. **e)** Detailed view of the Central-Coupler Elements (Arginine Sensor, Arginine Finger, substrate-coupling helix 1, Walker A and Walker B) for the subunit interfaces highlighted in (D) (Subunit A/Subunit B, Subunit E/Subunit F and Subunit F/Subunit A). The observed shifting between secondary structural elements ranges from 3 Å to 7 Å. **f)** Left: H-bond network established by different polar residues in the Central Coupler of the Subunit A/Subunit B interface in the ClpX hexamer in PDB 6PP7. The wildtype residue Q208 plays a major role to allow the intra-subunit B communication between its substrate-coupling helix 1 and its Arginine Finger which also then establish H-bond contacts with the Arginine Sensor in Subunit A. Right: The presence of the mutated A208 residue abolish both the intra-subunit and inter-subunit H-bond interactions, leading to significant movements of secondary-structural elements in the Arginine Finger, Arginine Sensor, Walker-A and Walker-B, mainly in the A/B and E/F and F/A subunit interfaces, as detailed in E. These structural rearrangements weaken the overall H-bond network of the central coupler.

Class-I resembles the substrate-free ClpXP reported PDB 8E91^27^ (RMSD = 0.83 Å, Supplementary Fig. 13a). As in that study, we find that the pore-2-loop of the subunit A^W^ appears in an ‘extended’ conformation blocking the potential entry of a polypeptide substrate (Fig. 5a). Moreover, all other relevant ClpX motifs (Box II, Walker-A, Walker-B, Arg-finger helix, substrate coupling helix-2, substrate coupling helix-1, Arg-finger or Arg307, sensor-I or Glu303, and sensor-II or Arg-370, pore-1-loop and pore-2-loop) in all subunits of WMWMWM overlap well with the corresponding motifs and subunits in PDB 8E91^27^ (Supplementary Fig. 13a-b). Overall, in the absence of a substrate, disruption of the central coupler in WMWMWM does not result in a structural change relative to wild-type ClpX. Furthermore, pore-1-loops and pore-2-loops, known to be involved in substrate recognition^12^, are seen in Class-I to adopt the same conformation as in PDB 8E91 (Supplementary Fig. 13b), suggesting that substrate recognition is not affected by Q208A. Altogether, the fact that we can obtain fully assembled ClpX hexamers and that Class-I (no substrate) is identical to a ClpXP complex with the wild-type central coupler also demonstrate that the single-chain construct allows full formation of the ring without oligomerization defects.

### Repositioning of Arg-finger, sensor-I, and sensor-II enables intersubunit communication

From all the previous cryo-EM studies^6,7,27–31^, Class-III closely resembles (RMSD = 1.92 Å) the substrate-bound ClpXP complex harboring a wild-type central coupler of PDB 6PP7^7^ (Fig. 5d, Supplementary Table 4). The central coupler elements in Subunits B, C and D overlap well between these two structures, even though subunits B and D harbor Q208A while subunit C is wild-type (Fig. 5d). However, differences were observed involving repositioning of different key protein motifs at the interfaces between the other subunits (Fig. 5d-e). In particular, in Class-III at the Subunit A/Subunit B interface, the Substrate-coupling Helix 1 and Arginine Finger Helix in subunit B, as well as the Walker-A, Walker-B, Arginine Sensor and the ADP nucleotide in the Subunit A, suffer displacements ranging from 2 Å to 5 Å (Fig. 5e), relative to their positioning in PDB 6PP7. These significant movements are also observed at the interfaces of Subunit E/Subunit F and Subunit F/Subunit A (Fig. 5e). The repositioning of these motifs suggests that some ‘re-shaping’ of the intermolecular interactions in the central coupler could be taking place in WMWMWM.

We then inspected in detail the central coupler of ClpX in PDB 6PP7 at Subunit A/Subunit B interface (Fig. 5f), and observed that the wild-type Gln208 (Q208) residue in the substrate-coupling helix 1 in Subunit B, plays a key role in establishing a H-bonding network that allow both intra and intersubunit communication with the Arginine Finger Helix (in subunit B) and Arginine Sensor (in subunit A), respectively (Fig. 5). Interestingly, extensive biochemical studies of ClpX have suggested that Arg307 (Arg-finger)^7,32–34^ and Glu303 (sensor-I) located in the Arginine Finger Helix^7^ may be involved in sensing the ATP status of the anticlockwise neighboring subunit, which agrees with our observations of the H-bond network formed in PDB 6PP7. Therefore, mutated Q208A residue in subunit B^M^, abolish the formation of the intra-subunit H-bonds with the Arginine Finger, causing the repositioning of these secondary structural elements and their key residues (sensor I and sensor II). This repositioning ultimately disrupts the intersubunit communication with the Arginine Sensor, Walker-A, Walker-B and ADP nucleotide located in Subunit A^W^ in the interphase A^W^/B^M^ (Fig. 5f).

### A new ClpXP structure with an extended conformation of pore-1-loop and pore-2-loop

Class-II does not resemble any published cryo-EM structure of ClpX. Unlike class-III, the pore-1-loop and pore-2-loop of the subunit A^W^ are in an “extended” conformation, oriented to interact with the polypeptide substrate (Fig. 5b). In contrast to our Class-II, the pore-1-loop of subunit A of PDB 6PP7^7^ is in a retracted conformation like in our Class-III, (pore-2-loop is not observed in PDB 6PP7). The nucleotide occupancy in Class-II is the same as in PDB 6PP7, where subunit A has an ADP and subunits B, C, D, E, and F harbor ATPs (Supplementary Fig. 10d). Additionally, small conformational changes of Arg-finger, sensor-I, and sensor-II are observed mainly at the interface A^W^/B^M^ and E^W^/F^M^ relative to those in PDB 6PP7 (Supplementary Fig. 13c). Also, unlike our Class-II, in which the pore-1-loop is in an extended conformation towards the substrate, in other conformations, such as PDBs 6PP5, 6PP6 and 6PP8^7^, 9C87, 9C88, and 8V9R^28^, and 6VFS and 6VFX^29^, the pore-1-loop is not extended, but the entire subunit A is displaced closer to the substrate to interact with it. Moreover, Class-II also differs in the nucleotide states found in the mentioned structures^7,28,29^ (Supplementary Table 4). Taken together, these observations indicate that, while Class-III resembles PDB 6PP7^7^, Class-II differs significantly from previously reported cryo-EM ClpXP structures with substrate (Supplementary Table 4), and therefore it represents a new conformational state (see discussion below).

## DISCUSSION

Fei *et al.*^7^ proposed that ClpX operates via the Probabilistic Anti-clockwise Long-Step (PA/LS) mechanism (Fig. 6a). Following ATP (_T_) hydrolysis (*) into ADP (_D_) and Pi, the top subunit A of the spiral moves anticlockwise down to the bottom position F, which accounts for the power stroke (→) (A*_T_B_T_C_T_D_T_E_T_F_D_ → B*_T_C_T_D_T_E_T_F_T_A_D_ →…, where the subunits are listed in order from top to bottom). Consequently, this process should be sensitive to strong mechanical barriers or high opposing forces. Furthermore, any mutation that affects the conformational changes involving subunits A and F will hinder the power-stroke.

**Fig. 6.**
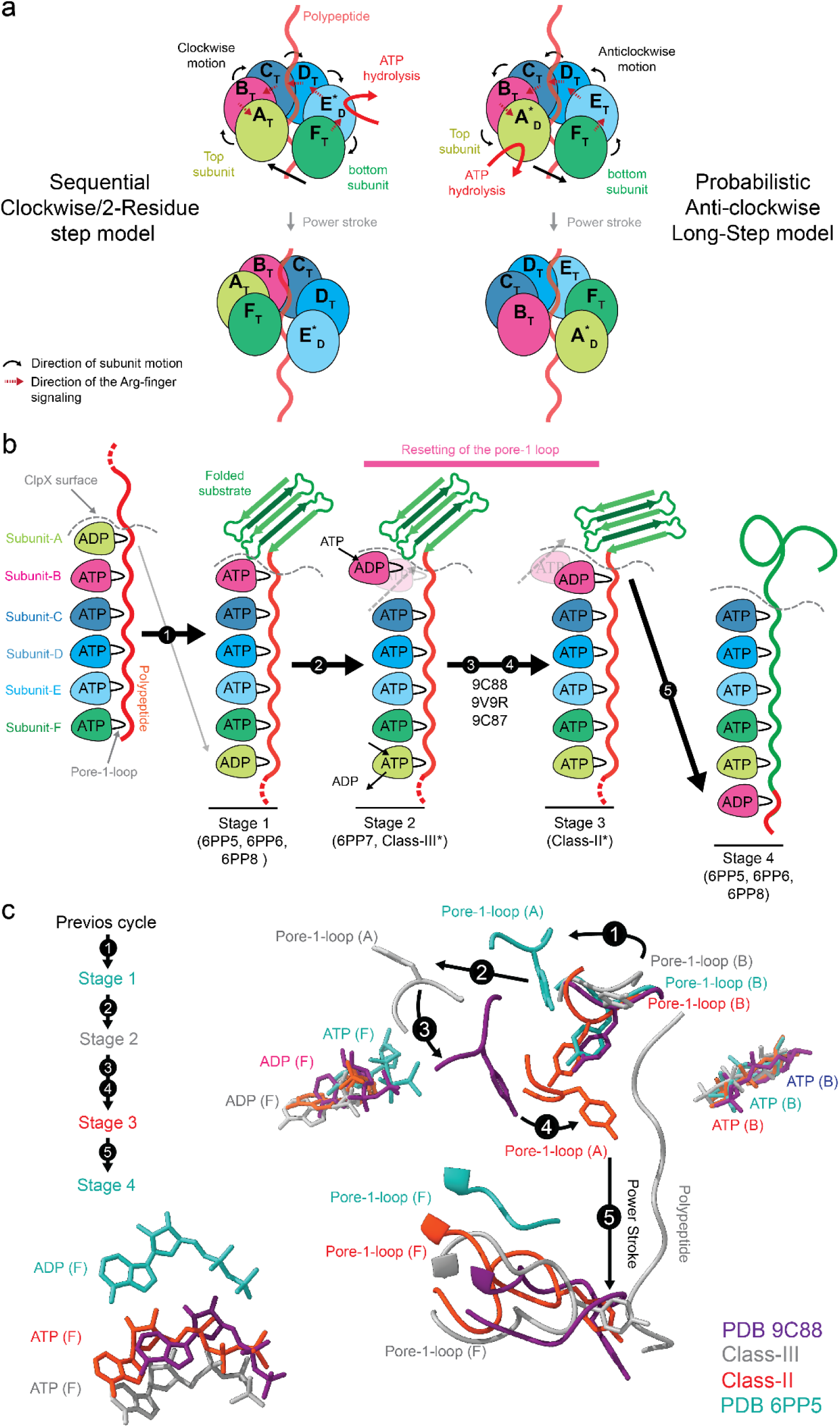
Model of the mechanism of action of ClpXP. **a)** Comparing the two contrasting models for the mechanism of power stroke generation in ClpXP; Sequential Clockwise/2-Residue Step model (SC/2R, left) vs Probabilistic Anticlockwise Long-Step (PA/LS, right). Each subunit is color-coded different. The subindex T or D means that a specific subunit is loaded with ATP or ADP, respectively. The red curved arrow indicates that subunit that hydrolyzes ATP. **b)** Schematic of the conformational states based on the cryo-EM structures observed previously^7^ and those described here. An additional state is observed between PDB 6PP5, 6PP6 6PP8^7^, Class-III* (*stage 4*) and 6PP7^7^ (*stage 2*) in the PA/LS model proposed by Sauer^7^. This intermediate, Class-II* (*stage 3*), is usually short-lived. *Refers to the coordinated version of Class-II and -III. Each subunit of the spiral (ABCDEF, from top to bottom) is represented with different colors. The folded protein (green color) is pulled down (power-stroke) by the downstream movement of top subunit A to bottom subunit F. ATP binding, hydrolysis (discontinuous arrows), and ADP release are annotated per ClpXP conformation. Between *stage 3* and *4*, the most recent cryo-EM structures (PDB 9C88, 9C87, 8V9R^28^) have a pore-1-loop in an intermediate conformation between its detached and extended states. **c)** Model of structural mechanism of the four conformational stages of the pore-1-loop to carry out the power stroke when the force generating subunit goes from position B to A and F in the spiral. Nucleotide status of subunit A, B and F of each structure are represented as sticks. Green = PDB 6PP5, Red = Class-II, Gray = Class-III, Purple = PDB 9C88.

On the contrary, in the Sequential Clockwise/2-Residue Step model (SC/2R), which was proposed as the general mechanism of action of AAA+ motors^1,35–37^, following ATP (_T_) hydrolysis (*), the subunit E moves clockwise down to the bottom position F, accounting for the power stroke, while the subunit in position F moves to position A (Fig. 6a). Accordingly, subunits A, B, C and D also move downwards becoming B, C, D, and E, respectively (A_T_B_T_C_T_D_T_E*_T_F_D_ → F_T_A_T_B_T_C_T_D*_T_E_D_). Note that in the SC/2R model, the top subunit A is never occupied with ADP, contrary to Fei’s Class 2 cryo-EM structure^7^. Instead, in the PA/LS model, ADP is located on the top subunit A, as observed in Fei’s Class 2 and our Class II and Class III structures. Also, in the SC/2R model, the downward motion of subunit E to F accounts for the power stroke. Thus, any mutation that would hinder the generation of force should affect the conformational changes associated with subunit E, which we do not observe (Fig. 5d).

Altogether, our findings strongly support the PA/LS model (Fig. 6b) and points to Class-II as the transient intermediate ready to perform the power stroke in ClpXP. First, Class-III (obtained with WMWMWM under ATP-RS conditions) has significant conformational changes in subunits A^W^ and F^M^ relative to the previously reported structure PDB 6PP7 (Class-2) while the other subunits (B^M^, C^W^, D^M^, and E^W^) remain unchanged (Fig. 5d). This result is predicted by the PA/LS model. Second, PDB 6VFS, 6VFX^29^, 6PP5, 6PP6 and 6PP8^7^ are post-power stroke conformations because the bottom subunit F has ADP (Supplementary Table 4) resulting from the transfer from subunit A to that position upon ATP hydrolysis and displaying the pore-1-loop extended to the substrate. Also, in the same structures, the subunit at the top position A is loaded with ATP having the pore-1-loop closer to the substrate but unable to do the power stroke. Third, PDB 6PP7^7^ represents a pre-power stroke state because subunits in position A and F are loaded with ADP and ATP, respectively, and the pore-1-loop in subunit A is not interacting with the substrate. Fourth, our Class-II has ADP and ATP bound to subunits A and F, respectively, like PDB 6PP7, but with the pore-1-loop of subunit A extended to interact with the polypeptide substrate. We interpret this structure as the transient intermediate between PDB 6PP7 and 6PP5 (Fig. 6b). Finally, the recent cryo-EM structures PDB 8V9R, 9C88 and 9C87^28^ are a combination of pre-and post-power stroke states because both top and bottom subunits of the spiral have ADP, and the pore-1-loop of the top subunit is not interacting with the substrate, whereas the bottom subunit shows this interaction.

We presume that the discovery of this new pre-power stroke conformation (Class-II) was enabled by the combination of (i) ATP hydrolysis inhibition via ATPγS, (ii) the hindering of force generation with Q208A in WMWMWM, and (iii) the presence of a mechanically stable substrate in front of the motor, making Class-II long-lived enough to be captured under cryo-EM. Indeed, without ATPγS, we did not observe Class-II in our cryo-EM analysis with Q208A because Class-II, although present, may still be too short-lived. This was also the case of previous structures hindering only ATP hydrolysis with the Walker-B mutation and/or in the presence of ATPγS^7,29^, or only challenging ClpX with an unfolding-resistant substrate^28^. Furthermore, WMWMWM is extremely sensitive to strong opposing force and high mechanical barriers (e.g. protein unfolding) but displays similar activity than WWWWWW in the absence of an external opposing force or when the barrier is minimal (translocation of unfolded polypeptides). These observations provide further support that the new conformation (Class-II) is on-pathway to the mechanism leading to the power stroke (Fig. 6b) and is only long-lived by the combination of the three conditions mentioned above.

Based on the cryo-EM ClpXP structures with substrate observed previously^7,28,29^ and those described here (all aligned relative to the invariant subunits C, D, E and substrate from PDB 9C88, PDB 6PP5, Class-II and Class-III), we propose a model in which the pore-1-loop of the top subunit in the spiral adopts conformations in four sequential stages during the power stroke (Fig. 6b, c). *Stage 1*: after power-stroke by subunit A, subunit B (loaded with ATP) occupies the position that subunit A had at the top of the spiral; see PDB 6PP5 (also 6PP6 and 6PP8^7^, Supplementary Fig. 14). Stage 2: the first of two ATP hydrolyses has taken place, the top subunit is found bound to ADP, and its pore-1-loop is in a *detached* conformation (the oxygen of Tyr153 OH-group is a distance δ = 26.3 Å away from the α-carbon of its closest substrate residue located within the ClpX pore, Supplementary Table 4 and Fig. 6c); see Class-III (and PDB 6PP7^7^). Stage 3: the previous ADP has been ejected, another ATP has been bound, and a second hydrolysis has taken place leading to the *extended* conformation of the pore-1-loop (δ = 5.4 Å closer to interact with the substrate in Class-II, Supplementary Table 4, Fig. 6c), followed by the power stroke. In the most recent cryo-EM structures (PDB 9C88, 8V9R, 9C87^28^, Supplementary Fig. 14), the pore-1-loop of the top subunit, occupied with ADP, is located at an intermediate distance between the *extended* (δ = 5.4 Å) and *detached* (δ = 26.3 Å) conformations. Thus, these structures are likely to represent pre-power stroke states with the pore-1-loop located *mid-way* (δ = 10.9 Å in PDB 9C88^28^, Supplementary Table 4, Fig. 6c) to interact with the polypeptide substrate (cyan color in Fig. 6c). Finally, Stage 4 or post-power stroke, when subunit F is loaded with ADP, located at the bottom of the spiral, and its pore-1-loop remains near (presumably interacting with) the substrate; see PDB 6PP5 (also 6PP6 and 6PP8^7^).

The extended and detached conformations of the pore-1-loop in the top subunit A observed in the present study, likely reflect the process of “resetting” that has been suggested for this loop^14,38^ (Fig. 6b). We propose that before an ATP hydrolysis is used during the power stroke, a previous ATP hydrolysis is required for this resetting. This model explains several basic features of the mechanism of action of ClpX. First, it rationalizes previous results indicating that ClpX consumes ∼2 ATP/power stroke with a step of 2 nm^9,14,16–19^. Second, it explains why mutations that slow down ATP hydrolysis (e.g. E185Q in the Walker-B motif^21^) or the presence of ATPγS^9,14,17^, make the dwell phase or the pause (i.e. dwells > 1 sec), during translocation, longer and more frequent in optical tweezers experiments. If hydrolysis of ATP is hindered, the motor remains steady for longer time before a translocation step. Third, it explains why WMWMWM hydrolyzes ATP significantly faster than the wild-type (Fig. 3e): *Stage 3* (Class-II) remains “stuck” right before the stepping because the mutation prevents subunit A from coupling its hydrolysis to a power stroke. As a result, subunit A undergoes several rounds of ATP binding and hydrolysis, consuming more ATP per translocation step than the wild-type hexamer. This same mechanism explains the observation that wild-type ClpX consumes more ATP/substrate when it is challenged with a client protein of greater stability^9,10,21,23,25,39–42^; *stage 3* is again stuck in front of the mechanical barrier.

Single-molecule and biochemical studies have revealed that (i) a wide variety of mutations of the pore-1-loop of ClpX alter the rate of ATP hydrolysis^14,15,38,43^, (ii) ATP hydrolysis rate per degraded substrate is correlated with the mechanical stability of the client protein^25,40–42^ and (iii) mutations in the Walker-B, Walker-A, Arginine sensor (R370) motifs that affect the ATP hydrolysis, and the presence of ATPγS slow down protein unfolding^9,10,21,23,39^. These observations suggest that a strong mechanochemical coupling within a ClpX subunit is necessary to generate sufficient force to overcome mechanical barriers. However, the mechanism underlying this mechanochemical coupling has remained hitherto unexplored. Our results provide direct evidence of the hypotheses proposed by Subramanian *et al*.,^20^. Indeed, the central coupler is the physical entity that (i) facilitates the communication between neighboring subunits and (ii) through which changes resulting from ATP hydrolysis quickly propagate from the ATP active site and to the pore-1-loop (mechanochemical coupling) for fast translocation and quick protein unfolding. Weakening the stiffness of the central coupler by the Q208A mutation, makes the mechanochemical coupling load-sensitive, so that a strong opposing force or a strong mechanical barrier such as a stable folded structure has the effect of *decoupling* the energy of nucleotide hydrolysis from the generation of force (playing the equivalent role of pressing the clutch in a combustion engine). As a result, in front of a strong load, the rate of ATP hydrolysis is “liberated” (de-coupled) from force generation and therefore it is no longer rate limited by the latter, as experimentally observed.

To summarize, we discovered a new pre-power stroke conformation in which the pore-1-loop is ready to generate force. The timing of the power stroke would depend on the speed at which the pore-1-loop of the top subunit A of the spiral receives the signal from its ATP site after hydrolysis via the central coupler. Furthermore, the ability of this subunit to do the power stroke depends only on its communication with its clockwise neighbor subunit B. Finally, our findings support a model in which one ATP is utilized during the pore-1-loop resetting and the second ATP during force generation. Therefore, the subunit occupying the top of the spiral (subunit A) plays a “special” role since the various adopted conformations of its pore-1-loop determine the generation of force during the power stroke, and the thermodynamic efficiency of the motor, both mediated by a rigid central coupler in subunits A and B. Because the central coupler is conserved^20^, our results are valid for other AAA+ enzymes.

## METHODS

### Purification of ClpX and protein substrates

All proteins (ClpX single-chain hexamers, substrates containing GFP and/or Titin) were purified by a similar workflow. Proteins were overexpressed in recA-deficient *E. coli* BL21 (DE3) cells^44^, and induced with 1 mM IPTG for 3 hours at 204 rpm when OD reached 0.6 – 0.8. For the case of single-chain ClpX constructs, induction was at 22°C at 204 rpm, but it was at 37°C in other cases. To prevent degradation, we used 1 mM of the serine protease inhibitor PMSF. For ClpP production, we did not use PMSF since ClpP is a serine protease. Cells were harvested at 4500 x g for 30 min at 4°C and resuspended in NiA buffer (20 mM HEPES pH. 7.6, 100 mM KCl, 400 mM NaCl, 20 mM imidazole, 10% glycerol). Lysis of 40 ml of resuspended cells was carried out using the homogenizer EmulsiFlex-C5 (AVESTIN Inc.) at 1100 bar for 15 min. After lysis, the solution was centrifuged at 16000 x g for 45 min at 4°C. Supernatant was passed through a 0.45 µm filter before putting in the 5ml HisTrap HP His tag protein purification column (Citiva). The 5ml HisTrap HP column was equilibrated with 5 volumes of NiA buffer with 10 mM βME, sample was loaded, washing was performed with 10 volumes of NiA buffer with 10 mM βME, and elution was carried out with 10 ml of NiB buffer (20 mM HEPES pH. 7.6, 100 mM KCl, 400 mM NaCl, 250 mM imidazole, 10% glycerol) with 10 mM βME. In all cases, flow rate was ∼3 ml/min at 4°C. Protein was monitored with Bradford (Biorad). We performed buffer exchange using the PD10 desalting columns (Citiva) with gel filtration buffer (50 mM Tris HCl pH. 7.5, 300 mM KCl, 0.1 mM EDTA, and 10% glycerol) with 1 mM DTT at 4°C. Only ClpX hexamers were further purified via size exclusion using a Superdex® 200 Increase 10/300 GL column (Citiva) in gel filtration buffer with 1 mM DTT at 0.2 ml/min at 4°C. For each ClpX hexamer construct, fractions were combined and concentrated up to ∼10 µM using 100K Amicon® Ultra 15 mL filter (Millipore Sigma). All proteins were stored in gel filtration buffer, aliquots were frozen with liquid nitrogen and stored at −80°C.

### Purification of ClpP

ClpP was induced and purified with HisTrap HP column as mentioned above. After elution with NiB with 10 mM βME, we performed buffer exchange using the PD10 desalting columns (Citiva) with Q50 buffer (50 mM Tris HCl pH. 8.0, 50 mM KCl, 10 mM MgCl2, and 10% glycerol) with 1 mM DTT. ClpP was further purified using a 5ml HiTrap Q HP anion exchange chromatography column. The HiTrap Q HP anion was equilibrated with 5 volumes of Q50 buffer with 1 mM DTT, sample was loaded, washing was performed with 5 volumes of Q50 buffer with 1 mM DTT, and gradient elution was carried out to 100% Q1000 buffer (50 mM Tris HCl pH. 8.0, 200 mM KCl, 10 mM MgCl2, and 10% glycerol) with 1 mM DTT in 20 min at 4°C. Fractions containing ClpP were combined and concentrated up to ∼50 µM using 30K Amicon® Ultra 15 mL filter (Millipore Sigma). We performed buffer exchange using the PD10 desalting columns (Citiva) with gel filtration buffer with 1 mM DTT at 4°C. ClpP aliquots were frozen with liquid nitrogen and stored at −80°C.

### Purification of TEV protease

TEV was purified using the pRK793 plasmid system in *E. coli* BL21 (DE3). IPTG induction, cell harvesting, and cell lysis were performed as mentioned above with some modifications. Before Lysis, cells were resuspended in Buffer A (50 mM Sodium Phosphate pH. 8.0, 500 mM NaCl, 40 mM imidazole, and 10% glycerol) with 10 mM βME at 4°C. TEV protease was purified using the HisTrap HP with Buffer A as the low imidazole solution, and gradient elution was carried out to 100% of buffer B (50 mM Tris-HCl pH. 8.0, 500 mM NaCl, 500 mM imidazole, and 10% glycerol) with 10 mM βME in 20 min at 4°C. Pure TEV protease was concentrated up to ∼30 µM with 10K Amicon® Ultra 15 mL filter (Millipore Sigma) with gel filtration buffer with 1 mM DTT. Final protein was aliquoted, frozen with liquid nitrogen, and stored at −80°C.

### Purification of Sortase 5M

The plasmid expression pET30b for sortase 5M was kindly provided by Prof. Andreas Martin. Sortase 5M was overexpressed in recA-deficient *E. coli* BL21 (DE3) cells^44^, and induced with 0.5 mM IPTG when OD was 0.5 – 0.7 for 4 hours at 30°C at 204 rpm. Cell harvesting and cell lysis was performed as mentioned above. Sortase 5M was initially purified with a 5ml HisTrap HP column (Citiva) using a gradient from Buffer A (50 mM Tris HCl pH. 7.5, 300 mM KCl, 0.1 mM EDTA, and 40 mM imidazole) to 100% of Buffer B (50 mM Tris HCl pH. 7.5, 300 mM KCl, 0.1 mM EDTA, and 1 M imidazole) in 20 min both with 10 mM βME at 4°C. Selected fractions were combined and dialyzed overnight with Buffer C (20 mM Potassium phosphate buffer pH. 6.0) with 1 mM βME with gently mixing at 4°C. Sortase 5M was further purified with a 5ml HiTrap SP cationic exchange column (Citiva) by gradient from buffer C to 100% of Buffer D (20 mM Potassium phosphate buffer pH. 6.0, and 250 mM KCl) for 20 min both with 10 mM βME at 4°C. Sortase 5M was finally purified via size exclusion with a HiPrep Sephacryl S-100 HR column using gel filtration buffer with 1 mM DTT at 0.3 ml/min at 4°C. Fractions containing pure Sortase 5M were combined and concentrated up to ∼800 µM using 10K Amicon® Ultra 15 mL filter (Millipore Sigma). Aliquots were frozen with liquid nitrogen and stored at −80°C.

### Biotinylation of ClpX hexamers

We designed a significantly more efficient labelling method of ClpX and single-molecule (SM) substrates from what was reported^14,16,17^. We C-terminally biotinylated 8 µM of *E. coli* ClpX single-chain hexamers with C-terminal sortag-(His)6-tag with 500 µM GGGK-biotin peptide (GenScript) and 10 µM sortase 5M mutant in the sortase reaction buffer (50 mM Tris HCl pH. 7.5, and 150 mM KCl) with 1 mM DTT at room temperature for 2 hours. Excess of GGGK-biotin peptide and sortase 5M were washed out via seven consecutive cycles of concentration and dilution at 6500 x g with gel filtration buffer plus 1 mM DTT at 4°C using 100K Amicon® Ultra 15 mL filter (Millipore Sigma). Biotinylation was confirmed via a retention assay with 2 µm streptavidin beads (Spherotech) in 5% PAGE native gel with 0.5X TBE buffer at 4°C. Biotinylation was further confirmed via standard Dot-blot assay in TBS buffer (20 mM Tris HCl pH. 7.5, 150 mM KCl) with Cy5-streptavidin over a nitrocellulose membrane with serial dilutions of biotinylated ClpX single-chain hexamers. Blocking solution was TBS buffer with 50 mg/ml BSA, excess of Cys-streptavidin was washed out with TBS buffer with 1 mg/ml BSA. All C-terminally biotinylated ClpX hexamers were diluted down to 250 nM, aliquoted, frozen with liquid nitrogen, and stored at −80°C.

### Chemical modification and Biotinylation of Single-molecule substrate

We designed a new SM substrate where we engineered a TEV cutting site (ENLYFQ//G) in the N-terminus followed by (Gly)_2_-GFP-(I27-Titin)_4_-(His)_6_-ssrA. Before labelling, the four I27-Titin moieties were chemically and permanently unfolded via carboxymethylation of the two cysteines buried^14,25^ with iodoacetic acid (Thermo ScientificTM) at a concentration equivalent to 100 times of the SM-substrate in the presence of 50 mM DTT and 2.5 M guanidinium chloride for 2 hours at room temperature with constant shaking and covered with aluminum foil. We performed buffer exchange of the carboxymethylated SM-substrate using PD10 desalting columns (Citiva) in TEV 1X buffer at 4°C. Permanent chemical unfolded Titins were confirmed with the red-shifted fluorescence spectrum from 315 nm to 345 nm of titin tryptophan^25^ using FELIXTM fluorimeter (Photon Technology Instruments).

The carboxymethylated SM-substrate was cleaved overnight with 10 µM of TEV protease in TEV reaction buffer (50 mM Tris HCl pH. 8.0, 0.5 mM EDTA) to generate the N-terminally (Gly)_3_ end, (Gly)_3_-GFP-(I27-Titin)_4_-(His)_6_-ssrA, for the sortase reaction. TEV-cleaved SM substrate was purified from the N-terminal fragment and TEV protease via size exclusion using the Superdex® 200 Increase 10/300 GL column (Citiva) in gel filtration buffer with 1 mM DTT at 0.2 ml/min at 4°C. We performed the N-terminal sortase reaction^45^ of ∼40 µM of (Gly)_3_-GFP-(I27-Titin)_4_-(His)_6_-ssrA with 125 µM of the peptide Biotin-MSDYDIPTENLPETGG in the presence of 10 µM sortase 5M in sortase reaction buffer for 2 hours at room temperature. The N-terminally biotinylated SM substrate, Biotin-GFP-(I27-TitinCM)_4_-(His)_6_-ssrA, was purified from the excess of Biotin-peptide and sortase 5M via size exclusion using the Superdex® 200 Increase 10/300 GL column (Citiva) in gel filtration buffer with 1 mM DTT at 0.2 ml/min at 4°C. Final protein sample was diluted down to 1 µM, aliquoted, frozen with liquid nitrogen, and stored at −80°C. Biotinylation was confirmed via a retention assay with 2 µm streptavidin beads (Spherotech) in 5% PAGE native gel with 0.5X TBE buffer at 4°C.

### Preparation of samples for optical tweezers

We mixed 750 pmol of C-terminally biotinylated ClpX hexamers with 2 µl of 0.14% BSA-passivated 2 µm streptavidin beads (Spherotech) and incubated for 5 min at room temperature. In the case of the N-terminally biotinylated SM substrate, we first ligated a 2 kbp-long dsDNA handle to 2 µm oligo-crosslinked polystyrene bead (OB, BangsLab) using T4 DNA ligase (New England Biolabs) in ligase reaction buffer for 1 hour at 22°C without inactivation. Half of the ligated handle-OB reaction was incubated with 2 pmol of neutravidin for 5 min at room temperature. Later, we added 1 pmol of N-terminally biotinylated SM substrate and mixed thoroughly for 5 min at room temperature. In both cases, ClpX-SB and OB-handle-neutravidin-SM substrate were finally diluted with 1 ml of filtered working buffer (ClpX 100) and injected into the microfluidic chamber.

### Optical tweezers data collection

All single-molecule trajectories were collected using a high-resolution dual-trap optical tweezers instrument with differential detection using a 1064-nm laser as previously described^46^, using a trap stiffness of ∼0.1 pN/nm. All activities were collected in ClpX 100 buffer (25 mM HEPES pH 7.6, 100 mM KCl, 0.5 mM EDTA and 20 mM MgCl_2_) with 5 mM ATP regeneration system^14,16,17^ to maintain the ATP concentration relatively constant during the experiment, 1 mM DTT, and 0.5 µM ClpP to ensure protein degradation^14,16,17^. The working buffer was passed through 0.2 µm filter. To prevent photodamage^46,47^, we also added 0.075 mg/mL Glucose Oxidase, 5 mg/mL Catalase, and 0.2% glucose as oxygen scavenging system to the working buffer just before usage. Activities were collected for ∼90 min to prevent significant effect of acidification on ClpX activity. Biotinylated ClpX hexamers and single-molecule substrates were immobilized separately on 2 μm-sized polystyrene beads held in two optical traps. After putting the two beads closer to each other, robust engagement of the ssrA-tag by ClpXP was observed by stable tether upon a slight increase of the force to 2-3 pN. Only tethers ranging between 600 – 700 nm size were selected to monitor activity because DNA handle attached to the single-molecule substrate is ∼2 kbp-long. At 2-3 pN and for ∼2-3 sec, we visually monitored the decrease of the bead-to-bead distance and the increase of force, which represent the active force generation and polypeptide translocation by ClpXP. Once we observed an active ClpX, the opposing force is automatically set to ∼6 pN, traps are fixed to a constant position (passive mode), and we started acquiring data at 2.5 kHz. During this screening process we always missed the initial ∼1 sec of every activity. In a typical single-molecule trace, before reaching GFP, the polypeptide translocation lasted ∼5-10 sec (depending on how fast to start recording the data). Traces that lasted only for 1 sec because the tether broke were discarded. The experiment ends when we observed a sudden tether break, either with or without unfolding event. In all cases, the opposing force increased monotonically. By the end of each activity, we estimated the stiffness and off-set of the current pair of beads, which we used for data calibration. The instrument was controlled using a custom software written in LabVIEW^46^. Before analysis, we visually separated activities with tether break in a single step and down to 0 pN to select for single molecules. Activities with a tether break in multiple steps and/or down to forces above 0 pN were discarded.

### Data analysis of single-molecule trajectories

Raw data of single ClpX activities was downsampled 10 times to 250 Hz. We used a custom software written in MATLAB based on the Kalafut-Visscher step finding algorithm^46,48^. The algorithm finds the steps by minimizing the quadratic error, Σ(data-fit)^2^, when a step is to be placed in the data. Each step divides the data into sections, and the height of the step is the mean value of the data in the section. The algorithm tests placing a step at every point in the data, calculates the resulting quadratic error, and chooses the step with the smallest error value. The algorithm continues until the quadratic error falls below a threshold, which is set by the Schwarz information criterion (SIC)^48^. We found that using this regular SIC overfits the data^46^, so we multiply the SIC by 2.5 (penalty). This value is arbitrary, but we find that no significant difference arises if the penalty fluctuates between 2 and 3 (see simulations, Supplementary Fig. 4, 5). After running the code, we proceed to do a visual inspection if most of the positive and negative steps of the trace were correct. We allowed less than 5% of incorrect identification to the eyes to proceed to differentiate dwells from pauses, steps from backtracking, and unfolding from translocation. Incorrect means two clear steps identified as one or one clear step identified as two. When more than 5% of events were misidentified (usually during translocation), we applied again the algorithm with a penalty value below or lower than 2.5 in changes of 0.1 till we achieved less than 5% of incorrect identification to the eyes. Below we provide the definition and details of analysis of each of the events in a single-molecule trajectory.

### Step

In a single-molecule trajectory in the optical tweezers experiment, steps are detected as shortening of the tether length by a distance between 1 – 6 nm during polypeptide translocation^9,14,16–19,38^ (Supplementary Fig. 5a). In this event, ClpX hydrolyzes ATP and releases the phosphate, triggering the power stroke of the motor by the motion of the ClpX pore loops^14,17^. For the case of the ClpX wild-type hexamer, the steps considered for analysis were the consecutive ones in 1 sec time frame, which correspond to ∼6 steps assuming ∼160 msec of an average dwell time (see definition below). For the case of ClpX hexamers harboring one or more mutant Q208A subunits, we considered traces displaying at least 6 steps (1 sec threshold is insufficient because mutant constructs have longer and more frequent pauses). Although unlikely, segments of single-molecule trajectories containing less than 6 consecutive steps were not considered to determine step distribution and mean value. In general, most of the steps for analysis came from more than 15 consecutive ones. For plotting the step distribution, we used a bin size of 1 nm because this value is the shortest size of the step detected in our data and previous studies^9,14,16–19,38^. The mean step size (Fig. 2h) is calculated as the simple average of all steps per condition and per ClpX hexamer construct.

To have an independent measurement of translocation steps that do not rely on step-finding algorithms during polypeptide translocation, we performed the pair-wise distribution (PWD) analysis^16,46,49^. Because ClpXP of WWWWWW can also perform backtracking and pauses during polypeptide translocation, we selected only regions of the trace without those events with a size spanning at least 20 nm. We obtained the PWD by taking the autocorrelation of traces filtered by a moving average filter of 30 points or 83 Hz (for the case of the wild type). Distributions of individual traces were graded by their periodicity, by integrating the power spectrum over the frequency range ±10% of the average value of the peak distribution and only the top 30% were selected for averaging^46,49^. We also performed a similar analysis for WWWWWM (filtered to ∼83 Hz) and WWWWMM (filtered to 50 Hz). For the other constructs (WWWMWM, WWWMMM, WWMMMM, and WMWMWM) the PWD analysis did not show the expected periodicity of around 2 nm because in those traces, backtracking and pauses events are more frequent.

### Dwell

When the tether length does not change in time during polypeptide translocation (also includes the after and before a backtracking event) and lasts less than 1 sec (explained below). In this event, ClpX is stationary because it releases the ADP (from previous ATP hydrolysis) and binds a new ATP to make a new translocation step^14^. For plotting the dwell time distribution (Supplementary Fig. 5b), we used a bin size of 30 msec which is 2.5-fold the shortest dwell time detected in our data (∼12 msec). The mean dwell time (Fig. 2h) is calculated as the simple average of all dwell times shorter than 1 sec per condition and per ClpX hexamer construct.

### Backtracking

We found that during polypeptide translocation of single-molecule traces, the tether experienced, although unlikely, a lengthening ranging between 1 to 5 nm in a single event or in a stepwise manner, followed by normal translocation (Fig. 2g, Supplementary Fig. 5a). Because they are of comparable size to translocation steps, we referred to them as backtracking. In contrast, events of large tether increase (>>5 nm) have a wide distribution of sizes and appear to be associated with a temporary loss of grip of ClpXP for its polypeptide substrate^16^. It is possible that the backtracking events represent short periods during which the motor can “walk backwards”, pulling away the polypeptide chain from the ClpP chamber without disengagement. We quantified the backtracking frequency as the number of those events divided by the total number of forward and backward stepping events in a polypeptide translocation region of a trace. Then, the backtracking probability can be computed as the average of the backtracking frequency values of all traces for each ClpX hexamer mutant (Fig. 2f). Each backtracking step was considered individually even if they happened consecutively in a backtracking event.

### Pause

Up to now, there is not a rigorous definition of a “pause” in the context of ClpX during the translocation of an unfolded polypeptide. We also do not currently understand the biological role of a pause during protein degradation. Nonetheless, previous studies distinguished a pause from a dwell using an arbitrary threshold of 1 sec^14,16,17^ or 2.5 sec^9,19^. Despite the arbitrariness of this value, it helped past single-molecule studies to find the number of ATPγS bound to ClpX to determine if one or more subunits are actively hydrolyzing ATP during a translocation step^14,17^. For consistency and comparative purposes, we consider a pause to be a dwell longer than 1 sec since we have used the same experimental conditions of those previous studies^14,16,17^. We measured pause frequency by dividing the number of pauses over the total number of events in a polypeptide translocation region of a trace. If a trace does not have pauses, this value is equal to zero. Later, we calculate the pause probability (Fig. 2g) as the simple average of the pause frequency values of all traces for each ClpX hexamer mutant.

### Unfolding

After the monotonic decrease of the tether length during polypeptide translocation, we observed that the activity stopped for a while before the tether increased cooperatively. Later, a second phase of monotonic decrease of tether length happened before the activity ended (Fig. 2b). These features, along with the size of the cooperative lengthening (Supplementary Fig. 8), were consistent with GFP unfolding by ClpXP in the optical tweezers at similar experimental conditions^14,16,17^. Also, we were able to determine the two intermediates in the unfolding pathway of GFP reported previously^14,17^. We also measured the “pause before GFP unfolding” that is equal to the average time between the end of the first polypeptide translocation and GFP unfolding, which represent the ability of ClpXP to unfold a protein client. For cases when we did not observe GFP unfolding, because either the tether broke too early or the “pause before GFP unfolding” was too long (> 5 min), we limited our analysis only for steps, dwell and backtracking during polypeptide translocation.

For the analysis of all events mentioned above (steps, dwell, backtracking, pause and unfolding), we only used the single-molecule traces collected between ∼6 to ∼9.5 pN because, in that range, the polypeptide translocation velocity remained unchanged in all ClpX mutants. The instantaneous velocity and force sensitivity of the motor were determined over all the range of forces excluding dwells larger than 1 sec which are interpreted as pauses.

### Validation of the Kalafut-Visscher step finding algorithm

To validate the Kalafut-Visscher step finding algorithm, we generated simulated translocation traces of ClpXP, applied the algorithm and compared the ground-truth vs the found steps under different conditions (Supplementary Fig. 3, 4). The simulated traces (Supplementary Fig. 3b) were obtained with the following parameters:

### Step Size

We tested both a 2 nm fixed step size (Supplementary Fig. 3c, d) and a 1, 2, and 3 nm equally distributed step size (Supplementary Fig. 3e, f), because we do not have the resolution to distinguish between these two possibilities.

### Step Time

Gamma distribution with shape factor 2. Two ATPs are burned per cycle with the release of two ADPs (one from the previous cycle after the power stroke and one for the resetting of the pore-1-loops in the present cycle), being ADP release rate-limiting^14^. Therefore, at least 2 rate-limiting events (shape factor 2) occur between every step. We used a mean dwell time of 0.2 sec to obtain the average experimental velocity of ∼10 nm/s.

### Sampling frequency

2.5kHz, which is the one we used in our data collection.

### Noise

The estimated noise of our data, by taking the standard deviation of (data – fit), is between 2.5 and 3.5 nm with the mode being ∼3 nm, with some as large as 4.5 nm (Supplementary Fig. 3a). Therefore, in our simulations we tested the scenario of 3 nm (Supplementary Fig. 3c-f) and the worst-case scenario of 4.5 nm (Supplementary Fig. 4a, b).

After obtaining the simulations, we applied the Kalafut-Visscher step finding algorithm with a penalty value of 2.5, because for most of the traces we used a value of 2.5 ± 0.2 for analysis. We also tested a penalty value of 2 and 3 to demonstrate the robustness of the algorithm (Supplementary Fig. 4c, d).

### Worm-like chain analysis of GFP unfolding

To determine if the unfolding events we observed corresponded indeed to the unraveling of GFP by ClpXP, we used the worm-like chain (WLC) model^50–52^ to determine the contour length of the full unfolding transitions as follows (Supplementary Fig. 8),

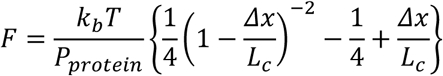

Where *F* is the unfolding force, *k_b_T* is the thermal energy, *Δx* is the change in extension during the unfolding event, *P_protein_* is the persistent length of the polypeptide, and *L_c_* is the contour length of the unfolded protein. This length can be converted to aminoacids using the value of 0.35 nm/aa^53^. The expected *L_c_* value during the full unfolding transition is calculated as the number of aminoacids contained in the β-barrel of GFP (∼217 aa or ∼76 nm) minus the initial distance between the two residues immediate and outside the folded GFP structure (∼2.6 nm). The subtraction gives the expected value of *L_c_* ∼73.4 nm.

Like previous studies of GFP unraveling by ClpXP^14,16,17^, unfolding events we observed happened without, with one or with two intermediates (Supplementary Fig. 8). Therefore, we considered the full length of the change in extension *Δx* regardless of unfolding happening via none, one or two intermediates. The corresponding force *F* considered in each trace was where the pause before unfolding happened. We then grouped all the data points (*Δx*, *F*) and used the WLC model to determine *L_c_* of the unraveled protein, using *P_protein_* = 0.65^14,16,17,53^ nm and *k_b_T* = 4.1 pN.nm at 25°C (Supplementary Fig. 8). We found *L_c_* to be 71.7 ± 0.9 nm (STD ∼ 10.8 nm) which is nearly the same as the theoretically expected value of ∼73.4 nm. Therefore, the 165 unfolding events used in our analysis (Fig. 4a-e), from the all the ClpX hexamer constructs tested in the optical tweezers with opposing force ranging between 6.5 and 9.5 pN, corresponded to the unraveling of GFP by ClpXP.

### Force sensitivity of single-chain ClpX hexamers

In passive mode, in our experimental conditions, each single-molecule activity starting at ∼6 pN went all the way to ∼9.5 pN. To cover a wider range of force, we repeated similar experiments starting with increment of ∼3.5 pN until we observed slowing down of the polypeptide translocation velocity. When the external force applied equals the maximum force generated by the motor, the velocity reaches zero value^26^. we fitted the force dependence of the pause-free velocity to a single-barrier Boltzmann equation^14,26^ (Fig. 4h),

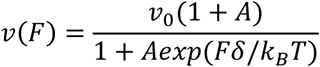

Where *k_B_* is the Boltzmann constant, *T* is the temperature, *v*_0_ is the velocity at zero force (nm/s), *δ* is the distance to the transition state (nm), and *A* is the dimensionless constant that determines the force sensitivity of the motor^14,26^. We estimated the external force at which the polypeptide translocation velocity is half of its maximum value F1/2^14,26^.

### Mant-ATP binding and SDS-PAGE degradation essays

We measured the equilibrium binding of single-chain ClpX wild-type and full Q208A mutant to the fluorescent ATP analog mant-ATP (2’-(or-3’)-O-(N-Methylanthraniloyl)adenosine 5’-triphosphate)^23^. Mant-ATP fluorescence increases upon binding to ClpX. Therefore, we used 1 µM of mant-ATP (Thermo ScientificTM) and titrated different ClpX concentrations (0, 0.05, 0.1, 0.5, 1, 2, 3, and 5 µM) in ClpX 100 buffer with 1 mM DTT lacking MgCl_2_ and in the presence of 5 mM EDTA to prevent ATP hydrolysis. We performed a scan emission from 400 to 500 nm using an excitation wavelength of 350 nm^23^ using FELIX^TM^ fluorimeter (Photon Technology Instruments). The binding curve is normalized value of the maximum absorbance intensity of mant-ATP per ClpX concentration relative to the maximum absorbance of the mant-ATP in the absence of ClpX vs each ClpX concentration tested. Each data point was triplicated (Supplementary Fig. 7b, inset).

We monitored the degradation activity of 14 µM Titin^CM^-ssrA by 0.6 µM single-chain ClpX hexamers in complex with 1.8 µM ClpP, 5 mM ATP regeneration system, and 1 mM DTT in ClpX 100 buffer at 22°C in 50 µl reaction (Fig. 2e). The reaction was initiated with Titin^CM^-ssrA, and took 7 µl sample at 0, 3, 5, 10, 15, 20, and 40 min incubation to mix with 4 µl of 4X SDS-loading buffer, heated at 95°C for 5 min, loaded in a pre-casted denaturing SDS-PAGE gel (Biorad), and run in Tris-Glycine buffer for 90 min at 22°C. we used the AcquaStain Protein Gel Stain (Bulldog Bio) to stain the gel. Titin^CM^-ssrA band intensity was quantified using ImageJ.

### Measurement of ATP hydrolysis rate

We determined the ATP hydrolysis rate of all ClpX hexamers tested in the optical tweezers using a coupled NADH-coupled assay (Supplementary Fig. 7). In this assay, the ADP, generated from ATP hydrolysis, with 2-phosphoenolpyruvate (PEP) are enzymatically converted by pyruvate kinase to ATP and pyruvic acid, which the lactate dehydrogenase (LDH) uses to convert NADH to NAD+ and lactate. Therefore, with this enzymatic assay we measured the rate of ATP hydrolysis as the oxidation rate of NADH to NAD+^54^. The NADH oxidation rate will be equivalent to the decrease rate of NADH absorbance at 340 nm, measured using a spectrophotometer in a 96-well plate^14,16,17^. The assay was carried out using three solutions. First, we freshly prepared 5X regeneration system (5X-RS) containing 15 U/ml PK (Sigma Millipore), 15 U/ml LDH (Sigma Millipore), 5 mM NADH (Sigma Millipore), and 37.5 mM PEP in 1X ClpX 100 buffer. Second, we prepared the ∼3X protein mixture containing 1.2 µM single-chain ClpX hexamer, 7.2 µM ClpP, 3 mM DTT, and 1.5 mM of TitinCM-ssrA in ClpX 100 buffer^14,16,17^. We used a permanently and fully titin-ssrA substrate at saturating concentration (∼500 µM) to ensure permanent polypeptide translocation by the ClpXP complex in solution which can be comparable to polypeptide translocation in our optical tweezers assay ^14^. Third, we freshly prepared the 5X initiation solution containing each ATP concentration in ClpX 100 buffer (5X-ATP): 25, 50, 100, 200, 300, 500, 750, 1000, 2000, 4000, 10000, and 50000 µM. We made the final 50 µl reaction volume with 14 µl of ClpX 100 buffer, 10 µl of 5X-RS, 16 µl of the 3X protein mixture, start the reaction with 10 µl of 5X-ATP, and mix thoroughly. We transferred the 50 µl reaction to a 96-well plate-reader and measured absorbance at 340 nm at 22°C for 40 min each 21 sec. The final ATP concentrations in the reaction were 5, 10, 20, 40, 60, 100, 150, 200, 400, 800, 2000, and 10000 µM. Each reaction at different ATP concentration was triplicated. The rate of ATP hydrolysis was calculated as follows,

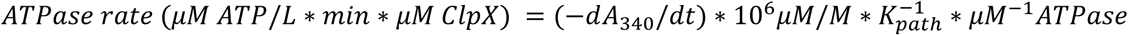

Where *dA*_340_/*dt* is the steady-state rate of NADH oxidation, and 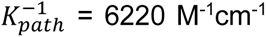 is the inverse of the molar absorption coefficient of NADH for a specific optical path length^20^.

### Assembly of ClpXP complexes for cryo-EM study

For the complex assembled in ATP conditions, we mixed 4 μM WAWAWA, 15 μM (ClpPx7)_2_, 5 mM ATP-RS, and 8 μM GFP-Ti^V15P,V13P^-(His)_6_-ssrA in 1X ClpX 100 buffer, and 1 mM DTT. Substrate was added at the end, mixed rapidly, incubated for 1 min at room temperature, and applied to the cryo-EM grids. Another sample was prepared adding also 200 μM of ATPγS before 1 min incubation.

### Cryo-EM sample preparation and data collection

Cryo-EM specimens were prepared on C-flat-1.2/1.3 400 mesh copper grids (Protochips) that were glow-discharged using a Tergeo-EM plasma cleaner (PIE Scientific). Then, 3.5 μl of sample were deposited on the grids, blotted for 10 sec with a blot force of 10 at 22°C in 100% humidity, and vitrified by plunging into liquid ethane using a Vitrobot Mark IV (Thermo Fisher Scientific). Electron micrographs were acquired as dose-fractionated movies in a 200 keV Talos Arctica cryo-electron microscope (Thermo Fisher Scientific) using a K3 direct electron detector (Gatan) operated in super-resolution counting mode. The microscope was set to 36,000X magnification (super-resolution pixel size of 0.5705 Å/pixel) with a total exposure dose of 50 electrons per Å^2^ fractionated across 50 frames. A total of 12,697 movies in ATP conditions and 25,584 movies in ATPγS conditions were collected with defocus values ranging from approximately –0.8 μm to – 2 μm. SerialEM^55^ was used to automatically control the data collection which parameters are detailed also in Supplementary Table 2. The cryo-EM data revealed the presence of full ClpXP complexes and free ClpP.

### Cryo-EM data processing

The processing of the cryo-EM data was performed using cryoSPARC v4.4.1^56^. Movie frames were aligned using the Patch Motion Correction program and binned 2x (to 1.141 Å/pixel). Then, defocus estimation and the Contrast Transfer Function (CTF) fitting were performed using the Patch CTF Estimation program. In the corrected micrographs, we can readily observe particles that correspond to the ClpXP complex as well as to the free ClpP (Supplementary Fig. 10a). Data processing was conducted separately for conditions in which the complex was assembled with ATP or ATPγS (Data Set 1 and Data Set 2 in Supplementary Data 1, respectively). A preliminary round of data analysis using a subset of each data set was performed, where the particles were picked using the blob-picker program to generate 2D classes that served for a follow template-based particle picking.

For Data Set 1, a subset of 5,861 particles was used to produce an ab initio 3D reconstruction. This volume was then used as a reference for a Heterogenous Refinement round (N = 3) on ∼700k particles selected from several rounds of 2D classification. From this Heterogenous Refinement round, one class of about 226k particles (∼35%) displayed a stable ClpXP complex and was subjected to Homogenous refinement. Then, we performed a global 3D classification without alignment of this population, using only a solvent mask auto-generated on-the-fly. From this classification, one population of ∼86k particles (∼39%) was identified with the substrate inside the ClpX hexamer while another class of ∼74k particles was observed to be in a substrate-free state. The population containing the substrate was selected, further cleaned to 76,882 particles and subjected then to non-uniform refinement. At high-threshold, this refined reconstruction showed a clear translocated substrate, while at low-thresholds a noisy ill-defined density appeared on top of the ClpX hexamer which was attributed to the non-translocated substrate region (Supplementary Fig. 10b). Then, a soft mask, involving only ClpXP including the translocated substrate within the ClpX hexamer, was used to perform particle subtraction followed by non-uniform refinement to finally produce the ClpXP class-III at 3.4 Å overall resolution (Supplementary Fig. 10, 11).

For Data Set 2, a similar workflow was performed (Supplementary Data 1). A subset of ∼245k particles was subjected to Homogeneous Refinement followed by a 3D classification without alignment. Here, two classes, accounting for ∼26% (∼63k particles) and ∼35% (∼59k particles) of the population, showed the best structural features, and were refined independently. The first class mentioned above (∼63k particles), was selected and its population further cleaned to ∼53k particles then subjected to non-uniform refinement. At high-threshold, this refined reconstruction showed a clear translocated substrate, while at low-thresholds a noisy ill-defined density appeared on top of the ClpX hexamer which was attributed to the non-translocated substrate region (Supplementary Fig. 10b). Then, a soft mask, involving only ClpXP with the translocated substrate within the ClpX hexamer, was used to perform particle subtraction followed by non-uniform refinement to finally produce the ClpXP class-II structure at 3.8 Å overall resolution (Supplementary Fig. 10, 11).

As mentioned before, in both data sets a particular 3D class corresponding to the substrate-free ClpXP complex was identified (denoted here as ^(A)^ and ^(B)^Substrate-free ClpXP complexes in Supplementary Data 1). The particles were merged and further cleaned, resulting in ∼68k particles producing the ClpXP Class-I structure obtained at 3.7 Å global resolution (Supplementary Fig. 10, 11). All Q-scores can be found in Supplementary Table 3.

### Model Building and Refinement

The final refined maps of ClpXP Class-I, -II and -III, were sharpened with B-factors of −85, −30 and −30, respectively, to prepare them for the initial model building. For Class-I, the initial model was obtained by rigid-body fitting the atomic coordinates of the substrate-free wild-type ClpXP complex, PDB 8E91^27^, into the corresponding sharpened map using UCSF ChimeraX^57^. For Class-II and Class-III complexes, the starting coordinate model was also obtained from PDB 8E91^27^, but the pore-1 and −2 loops for subunit A, as well as the substrate bound, were rebuilt manually. For each complex, the models were then iteratively rebuilt in COOT^58^ and refined using the real space refinement program in PHENIX^59^. For Class-I, the final sharpened cryo-EM map was obtained using the DeepEMhancer sharpening program^60^. For Class-II and Class-III, the final sharpened maps were obtained using the LocScale Program^61^. All validation and refinement statistics are shown in Supplementary Table 2.

### Data Statistics

All single-molecule and cryo-EM datasets were independently reproduced across multiple rounds of data collection. To minimize potential systematic errors, data for different experimental conditions were interleaved and collected in a randomized order. No statistical method was used to predetermine sample size; instead, they were established based on previous publications and data collection was gathered until the standard error of the mean fell below 10% of the average. Cryo-EM sample size was determined by collecting data until the global resolution was below 4 Å. All sample sizes were sufficient for demonstrating statistically significant differences when relevant. Predefined data exclusion criteria were applied to the single-molecule and cryo-EM datasets (all details are mentioned in the material and methods of each procedure). The Investigators were not blinded during experiments and outcome assessment. However, potential bias was mitigated by analyzing all data sets via an identical, automated, and predefined computational pipeline.

## DATA AVAILABILITY

The single-molecule data collected in this study has been deposited in the Zenodo database under the accession code 21879510 (https://zenodo.org/records/21879510). The cryo-EM density maps and their respective atomic coordinate files have been deposited to the Electron Microscopy Data Bank (EMDB) and Protein Data Bank (PDB) under the following accession codes: EMD-70686 and PDB 9OPA (ClpXP Class-I), EMD-70772 and PDB 9ORD (ClpXP Class-II), EMD-70861 and PDB 9OTX (ClpXP Class-III).

## CODE AVAILABILITY

The LabVIEW and MATLAB code used in this study has been deposited in the Zenodo database under the accession code 4632595 (https://doi.org/10.5281/zenodo.4632595) and 18810738 (https://zenodo.org/records/18810738), respectively.

## ACKNOWLEDGEMENTS

We thank the assistance of Graeme Fenton, Dilara Zenel, Vijay Prathigudupu, Ledet Getachew, and Jaquie Ho during various stages of the biochemistry, cloning of all proteins, and collection of optical tweezers data. We thank Eva Nogales for providing her laboratory facilities. We appreciate the valuable discussions with and feedback from members of the Kuriyan, Nogales and Bustamante Lab. We also thank D. Toso and R. Thakkar at the Cal-Cryo EM facility at QB3-Berkeley for helping with cryo-EM data acquisition, and P. Tobias and K. Stine for computing support.

## AUTHOR CONTRIBUTIONS

R.S. Cloned/produced all proteins and developed the single-molecule assay. A.T. Developed the software for collection/analysis of optical tweezers data. R.S., J.Km. Collected optical tweezers data. A.F. Prepared cryo-EM samples, collected and analyzed the cryo-EM data. R.S., Z.K. Developed the Biochemistry. R.S., J.Km., A.L. Performed the Biochemistry. R.S., J.K., C.B. guided the project, R.S., A.F., C.B. wrote the manuscript.

