## Supplementary Material for "The Central Coupler of ClpXP Controls Intersubunit Communication and Couples Intrasubunit Chemical Energy with Force Generation"


**This file contains**

Supplementary Results and Discussion
Supplementary Figures 1 – 15
Supplementary Tables 1 – 5

### SUPPLEMENTARY RESULTS AND DISCUSSION

#### Validation of the step finding algorithm with simulations

For the 2nm fixed step size, we see a great performance of the step finding algorithm because it gives the same result as the ground truth (Supplementary Fig. 3c), with only very rare events of missing steps (a very small fraction of 4 nm in the step size histogram of Supplementary Fig. 3d). In the variable step size case (Supplementary Fig. 3e), the step finding algorithm has some trouble identifying 1 nm steps (the triplet of peaks of 1, 2, and 3nm step sizes should be equal area, but the 1 nm one is smaller). This under representation leads to on average longer dwells due to these missed steps (Supplementary Fig. 3f). When we performed the simulations using a 4.5 nm noise (Supplementary Fig. 4a), the population of missed steps (4 nm) increases in the step size distribution histogram (Supplementary Fig. 4b), but still a good performance of the algorithm. Finally, we do not observe a significant difference in the dwell time and step size distribution if we use a penalty value of 2, 2.5 and 3 in the analysis of simulated traces with the step finding algorithm (Supplementary Fig. 4c, d).

#### Efficient mechanochemical coupling requires appropriate intersubunit communication

The ATP burning rate of each subunit in ClpX has two components: (i) the rate ($k_s$) of signaling (⟵) received from its clockwise neighbor to hydrolyze ATP, which depends on the integrity of the latter's central coupler, and (ii) the rate at which the subunit hydrolyzes ATP ($k_h$), which depends, in turn, on the integrity of its own central coupler. $k_s$ is normal or slow depending on whether the clockwise neighbor subunit is W or M, respectively. $k_h$ is normal if the hydrolyzing subunit is W, effectively coupling the ATP hydrolysis with the power stroke, or fast if the subunit is M (mutation Q208A in the central coupler) and its ATP hydrolysis is decoupled (futile), not limited by the generation of a power stroke. For WMWMWM (Supplementary Fig. 15a), the ATPase activity is interlaced between three events of slow signaling followed by normal ATP hydrolysis (normal $k_h$ ⟵ slow $k_s$) for each W⟵M pair, resulting in a slow and effective ATP burning, and three events of normal signaling followed by fast ATP hydrolysis (fast $k_h$ ⟵ normal $k_s$) for M⟵W pairs, resulting in fast and futile ATP burning. In contrast, for WWWMMM (Supplementary Fig. 15b), the first and the second W⟵W pairs will undergo a normal and effective ATP burning (normal $k_h$ ⟵ normal $k_s$); the third W⟵M pair displays slow signaling followed by normal ATP hydrolysis (normal $k_h$ ⟵ slow $k_s$) resulting in a slow and effective ATP burning; the fourth and the fifth M⟵M pairs display slow signaling followed by fast ATP hydrolysis (fast $k_h$ ⟵ slow $k_s$) resulting in slow and futile ATP burning. Finally, the last M⟵W pair displays normal signaling following by fast ATP hydrolysis (fast $k_h$ ⟵ normal $k_s$) resulting in fast and futile ATP burning. As a result, WMWMWM has three slow effective and three fast futile ATP burning events, whereas WWWMMM has two normal effective, one slow effective, two slow futile, and only one fast futile ATP burning events (Supplementary Table 5). Accordingly, WMWMWM has a faster ATPase rate (Fig. 3a) and a larger consumption of ATP per step (Fig. 3d) than WWWMMM. This analysis also explains why WWMMMM has a slower ATPase rate than WMWMWM (Fig. 3a), since it has only one fast futile (M ⟵ W), one normal effective (W⟵W), one slow effective (W ⟵ M), and three slow futile (M⟵M) ATP burning events (Supplementary Table 5). Finally, the full wild-type, WWWWWW has all the W⟵W pairs with normal and effective ATP burning events, and therefore, the smallest ATPase rate and consumption of ATP/step (due to the highest mechanochemical coupling) among all constructs (Fig. 3a).

### SUPPLEMENTARY FIGURES AND TABLES

**Supplementary Figure 1.** Size exclusion Chromatogram of wild-type and Q208A ClpX monomers, and wild-type and full Q208A hexamer mutant. Inset: SDS-PAGE gel of the degradation of titinCM-ssrA by ClpXP in an interval of 40 min. The two peaks at ~19 ml and ~20 ml both have absorbance at 260 nm, and the latter coincided with free ATP. This result indicates that ClpX monomers carrying the Q208A mutation, can bind and retain the bound nucleotides for a considerable amount of time for ligands to be co-purified with the monomers during size exclusion chromatography.

**Supplementary Figure 2. Step finding analysis of a single-molecule trajectory of substrate degradation by ClpXP.** *Above*, representative single-molecule trace of ClpXP during polypeptide translocation and protein unfolding of GFP. The step finding algorithm can identify almost perfectly steps and the unfolding events and its intermediates. *Below*, close-up view of the quality of the determination of steps.

**Supplementary Figure 3. Validation of the step finding algorithm with simulations, effect of step size. a)** Noise distribution of the regions of polypeptide translocation without pauses and backtracking events from single molecule traces of the WWWWWW construct (n = 52 traces). **b)** Simulated trace (grey color) with the parameters of fixed step size = 2 nm and noise = 3 nm. For comparison, three examples of real traces are shown in different colors. **c)** Representative example of simulated trace at 2.5 kHz (light grey) and downsampled to 250 Hz (dark grey) for the case of step size = 2 nm and noise = 3 nm. The results of the step finding algorithm are shown in blue color (off-set in Y for clarity) vs the ground truth (black). **d)** Dwell time and step size distributions obtained with the step finding algorithm with a penalty value of 2.5 from simulated traces using a step size of 2 nm and noise of 3 nm. **e)** Representative example of simulated trace at 2.5 kHz (light grey) and downsampled to 250 Hz (dark grey) for the case of step size = 1, 2 and 3 nm and noise = 3 nm. The results of the step finding algorithm are shown in blue color (off-set in Y for clarity) vs the ground truth (black). **f)** Dwell time and step size distributions obtained with the step finding algorithm with a penalty value of 2.5 from simulated traces using and step size of 1, 2 and 3 nm and noise of 3 nm.

**Supplementary Figure 4. Validation of the step finding algorithm with simulations, effect of noise and penalty. a)** Representative example of simulated trace at 2.5 kHz (light grey) and downsampled to 250 Hz (dark grey) for the case of step size = 2 nm and noise = 4.5 nm. The results of the step finding algorithm are shown in blue color (off-set in Y for clarity) vs the ground truth (black). **b)** Dwell time and step size distributions obtained with the step finding algorithm with a penalty value of 2.5 from simulated traces using a step size of 2 nm and noise of 4.5 nm. **c)** Representative example of simulated trace at 2.5 kHz (light grey) and downsampled to 250 Hz (dark grey) for the case of step size = 2 nm and noise = 3 nm. The results of the step finding algorithm with penalty values of 2, 2.5 and 3 are shown in color blue, red and yellow, respectively (different off-set in Y for each fitting for clarity) vs the ground truth (black). **d)** Dwell time and step size distributions obtained with the step finding algorithm with a penalty value of 2, 2.5 and 3 from simulated traces using a step size of 2 nm and noise of 3 nm.

**Supplementary Figure 5. Step and dwell time distribution of constructs studied in the optical tweezers. a)** Distribution of Step size made by ClpXP during polypeptide translocation.

Steps with negative values correspond to backtracking events. WWWWWW (n = 2909 events), WWWWWM (n = 3162), WWWWMM (n = 3308), WWWMWM (n = 2166), WWWMMM (n = 2334), WMWMWM (n = 1437), and WWMMMM (n = 1809). **b)** Distribution of all dwell times during polypeptide translocation. Pauses (dwells > 1 sec) were excluded. WWWWWW, WWWWWM, WWWWMM, WWWMWM, WWWMMM, WMWMWM, and WWMMMM.

**Supplementary Figure 6. Pair-wise distribution (PWD) analysis.** PWD analysis of section of the polypeptide translocation traces selected, from the raw data, that do not contain visible pauses nor backtracking events. Not all traces for a given construct used for the step finding algorithm fulfilled such conditions, because we selected only a minimum distance of translocated polypeptide of 20 nm (around 10 steps). The more mutant subunits are present in the ClpX hexamer construct, the less likely it is to find such sections, the easier it is to miss such events and include them in the PWD analysis, and consequently, the periodicity gets lost. Number of traces used per mutant are indicated in each PWD plot."

**Supplementary Figure 7. ATPase rate at different ATP concentrations for different single-chain ClpX mutants. a)** ATPase rate vs ATP concentration for WWWWWW (blue), WMMMMM (Fuchsia), and MMMMMM (green) constructs. *Inset*, Binding curve of mant-ATP association to WWWWWW (blue) or to MMMMMM (green). **b)** ATPase rate vs ATP concentration for all the constructs tested in the optical tweezers besides WWWWWW, WWWWWM (red), WWWWMM (cyan), WWWMWM (grey), WWWMMM (purple), WWWWWW (wheat), and WWMMMM (orange). Each condition was triplicated. Error bar is the SEM.

**Supplementary Figure 8. Analysis of GFP unfolding. a)** Different types of unfolding trajectories by ClpXP via none, one (either I1 or I2), or two intermediates (I1 and I2). Traces are downsampled six-fold for clear visualization (~400 Hz). **b)** Analysis of the total change in extension for certain intermediate or the length of the full transition (nm) vs the unfolding force (pN) during the experiment. Circles are experimental data; dotted line is the fitting to the worm-like chain (WLC) model of polymer statistics (see methods for details). The type of transition and the number of events is indicated in each plot. **c)** Molecular model of the stepwise unfolding of GFP via intermediates I1 and I2. First, the expected contour length of the extraction of β11 from fully folded to I1 (Δβ11) intermediate is ~9.8 nm, which is close to estimated value of 7.3 nm. Second, the unfolding of β10-7 from I1 to I2 (Δβ11-7) has a theoretical contour length value of ~29 nm, being the experimentally measured ~27 nm. Finally, from I2 to fully GFP unfolded, we estimated ~35 nm of contour length, which is like the expected value of ~38 nm. For the GFP structure, β11 is shown in blue color, β10-7 are depicted in orange, and the rest of the protein in gray color.

**Supplementary Figure 9. Effect of a single Q208A mutation on the ability of ClpXP to disrupt the biotin-neutravidin interaction. a)** Representative extension (nm) vs time (sec) traces showing events before tether break and after translocation of the unfolded GFP polypeptide for the case of WWWWWW (top) and WWWWWM (bottom). We rationalized this event as the successful extraction of biotin bond to neutravidin by ClpXP. Therefore, we call this event 'Biotin Extraction time'. *Inset*, zoom in of hopping transitions for the case of WWWWWM. **b)** Inverse of the cumulative density function plot of the Biotin Extraction time of WWWWWW (grey) and WWWWWM (cyan).

**Supplementary Figure 10. Cryo-EM Analysis of ClpXP complexes of WMWMWM. a)** Left: A representative cryo-EM micrograph section. Examples full ClpXP complexes and free ClpP particles are indicated by white and red arrows, respectively. Right: Representative 2D class-averages showing both ClpXP complexes (left) and free ClpP (right). **b)** Cross-sections of intermediate cryo-EM maps (See Data S1, red squares highlighted) obtained for Class-I, -II and -III ClpXP complexes at different threshold levels. Class-I does not harbor a substrate bound. In class-II and -III, the EM density corresponding to the translocated and non-translocated substrate regions, is observed at different threshold levels. **c)** Fourier-Shell-Correlation (FSC) map vs map and map vs model (left), and Local-resolution colored density map (right) for the ClpXP Class-I, -II and -III. **d)** Close-up view of the Q208 wildtype or A208 mutated residue (top row), and the nucleotide bound (ATP or ADP, bottom row) for each ClpX subunit for three classes. **e)** Representative regions of the atomic coordinate model fitted into the cryo-EM density map for the three ClpXP classes.

**Supplementary Figure 11. Angular distribution and Directional Resolution Plots. a)** Angular distribution plots for the cryo-EM ClpXP class-I, -II and -III structures, as indicated. **b)** Directional Resolution FSC summary plots for the cryo-EM ClpXP class-I, -II and -III structures, as indicated. The reported sphericity score corresponds to the conical Fourier anisotropy ratio (cFAR).

**Supplementary Figure 12. Comparing the substrate within the ClpX pore for different structures.** The PDB codes of the structures are indicated in each square. The polypeptide substrate within the ClpX pore is depicted in gray color. The pore-1-loop (aminoacids 152 to 153, with the Tyr153 in sticks) of each subunit (from bottom to top) are color-coded as in figure 5.

**Supplementary Figure 13. Comparing cryo-EM ClpXP structures without substrate. a)** Overlapping of the ClpX coordinates of Class-I vs PDB 8E91. Bottom: close-up view of the overlapped structures in the Subunit E/Subunit F interface. The ClpX atomic model in Class-I (harboring the triple alternate mutation Q208A), closely matches the ClpX model in the substrate-free wild-type ClpXP structure PDB 8E91. **b)** Comparing the pore-1-loop (top) and pore-2-loop (bottom) of Class-I and PDB 8E91. **c)** Top: Overlapping of ClpX coordinates in Class-II vs PDB 6PP7. Bottom: close-up view of the overlapped structures in the Subunit E/Subunit F interface. The ClpX atomic model in Class-II (harboring the triple alternate mutation Q208A), shows significant conformational changes relative to the wild-type ClpX atomic model in PDB 6PP7. The major differences occurred along domains that 'sense' the nucleotide binding and hydrolysis.

**Supplementary Figure 14. Comparing the top, second top, and bottom subunits of other cryo-EM ClpXP structures with substrate. a)** for Fei *et al.,* 2020[1] and **b)** Ghanbarpour *et al.*, 2025[2]. Each PDB structure is depicted in different colors.

**Supplementary Figure 15. Model for explaining the higher ATP hydrolysis rate of WMWMWM relative to WWWMMM**.
**(a)** Model for WMWMWM
**(b)** Model for WWWMMM.

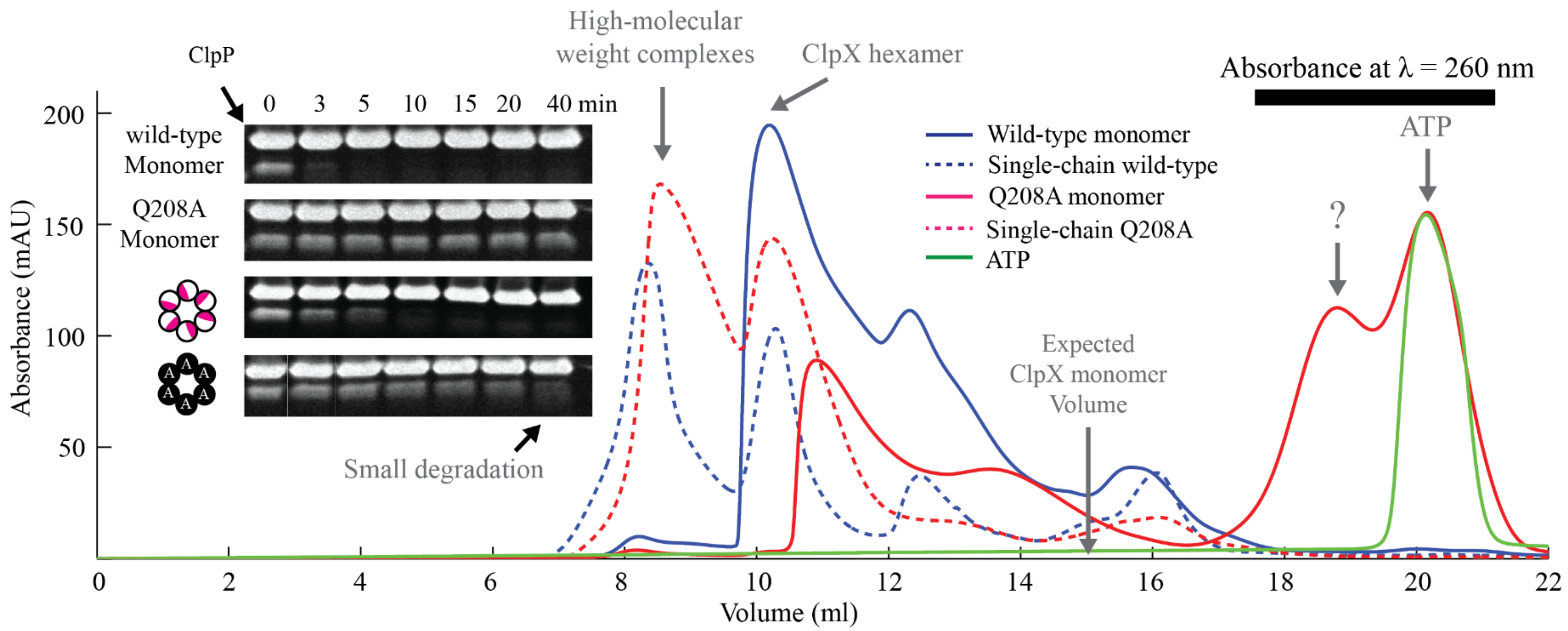


**Supplementary Figure 1**

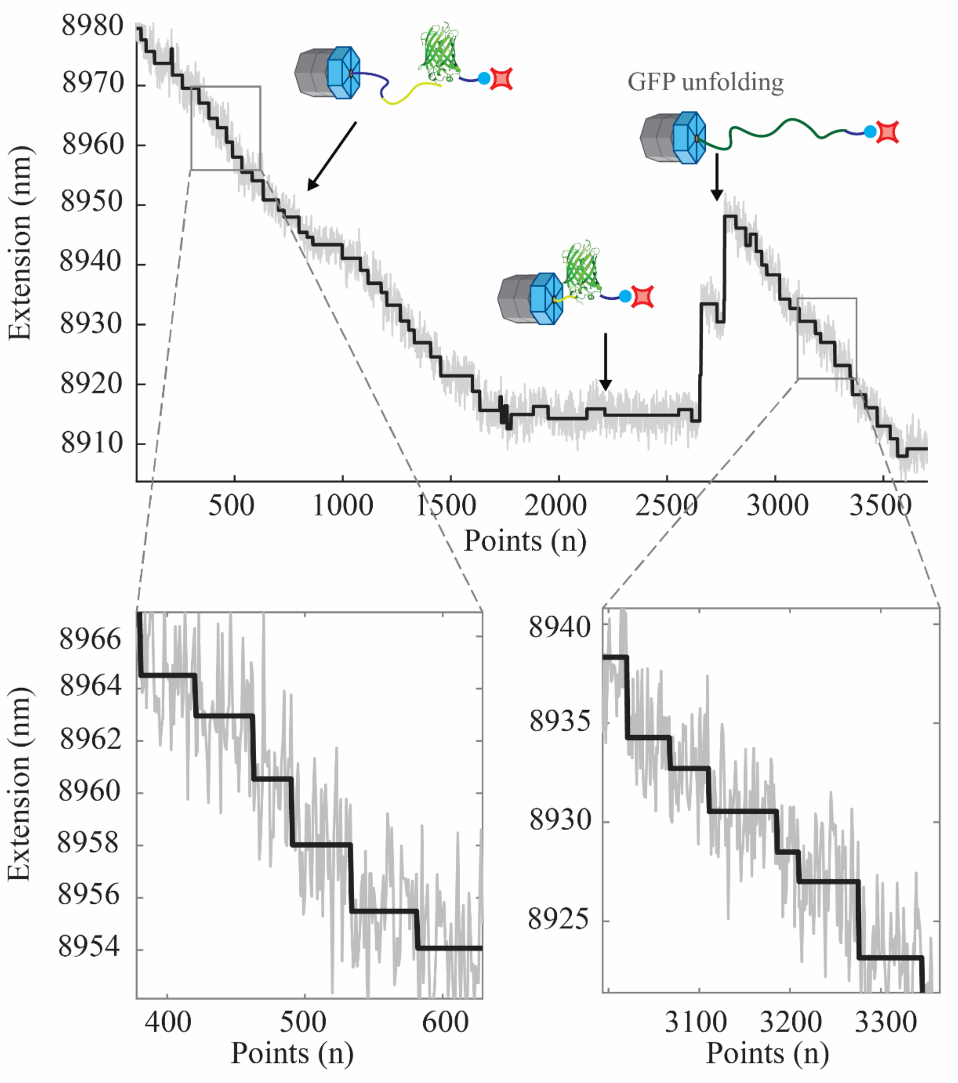


**Supplementary Figure 2**

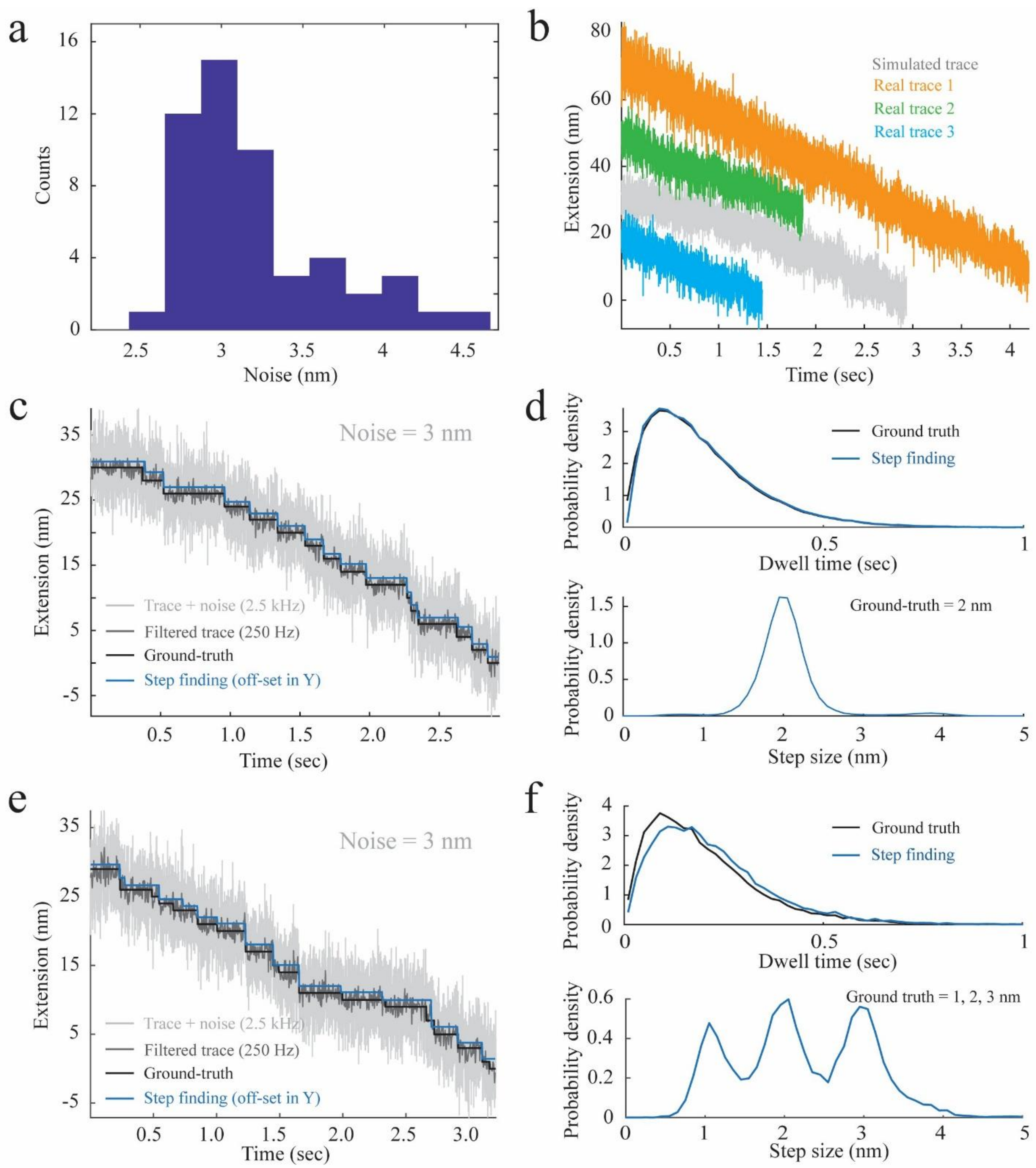


**Supplementary Figure 3**

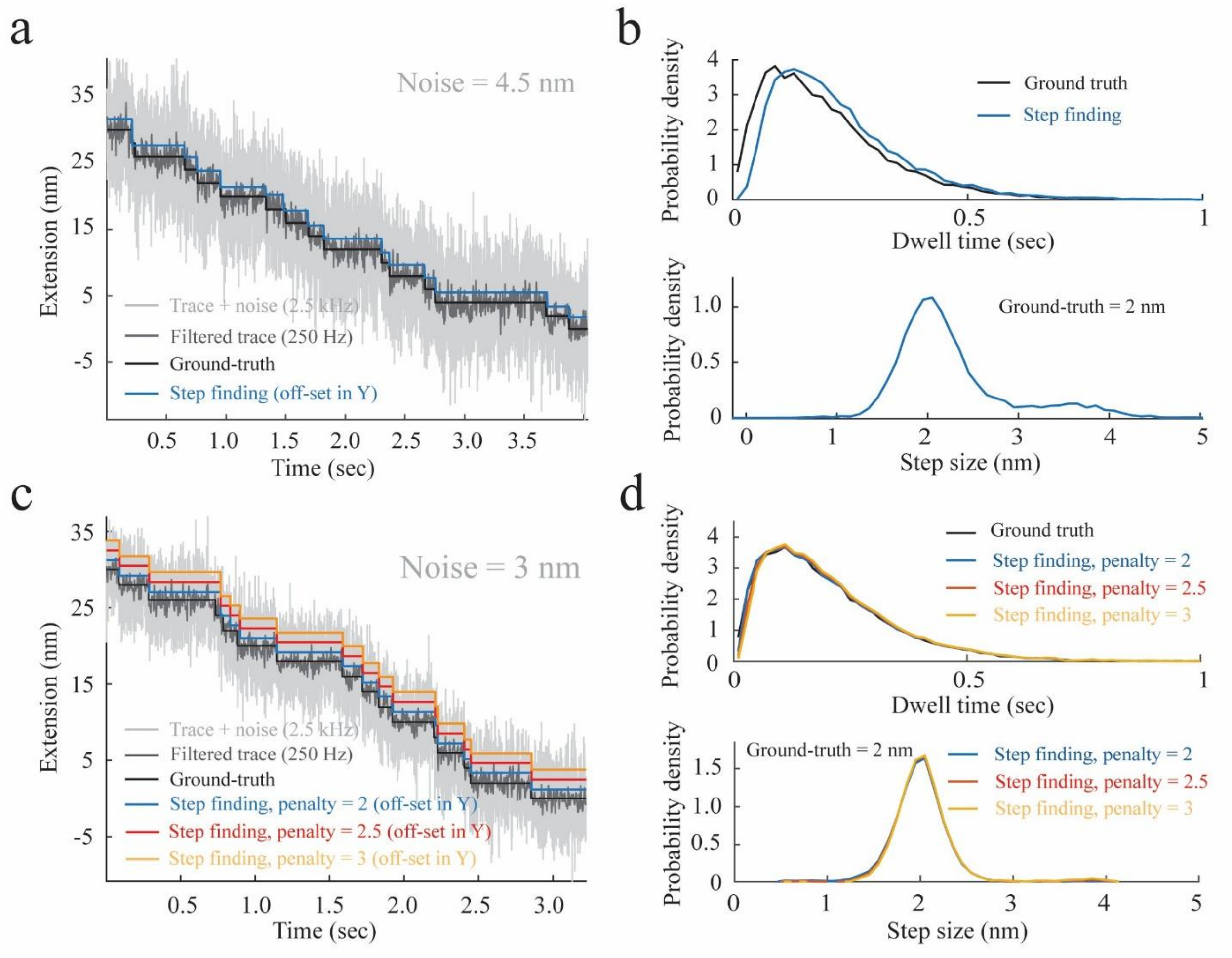


**Supplementary Figure 4**

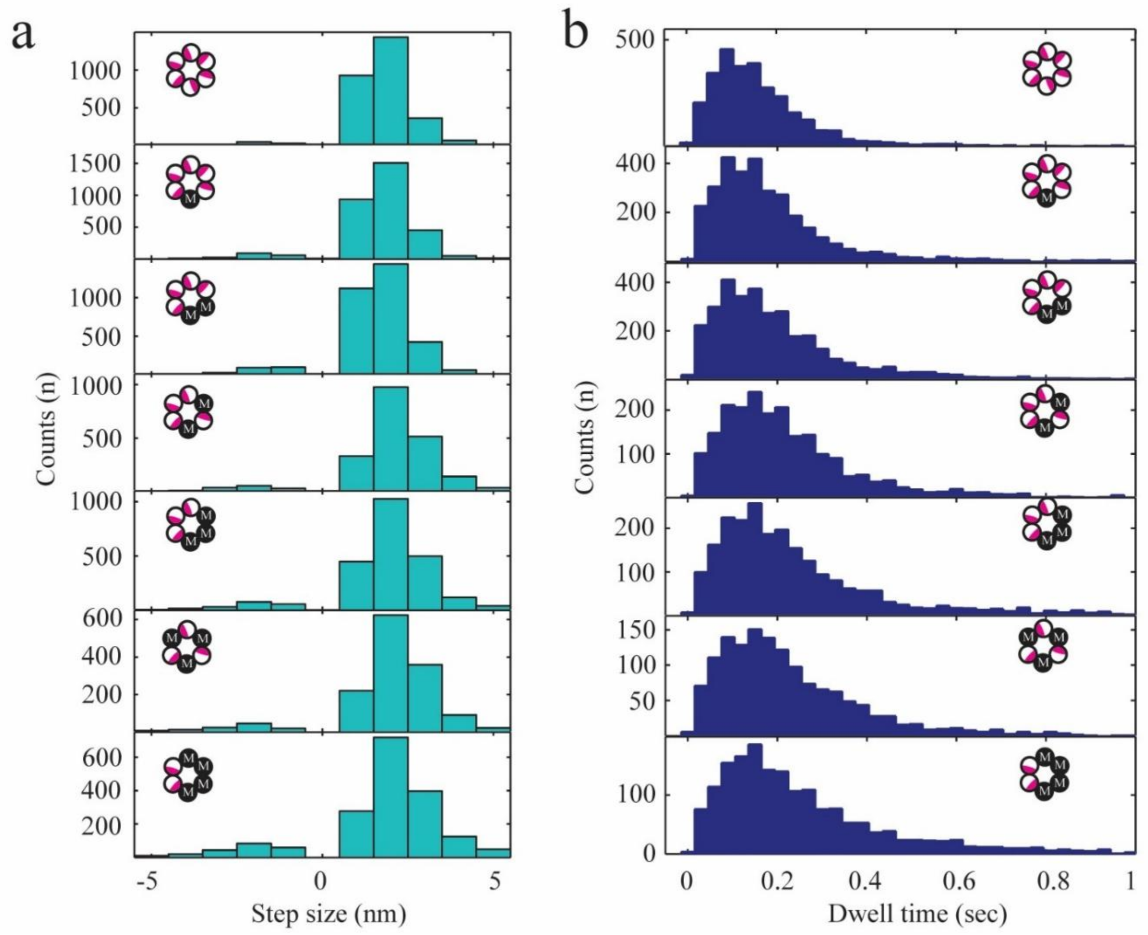


**Supplementary Figure 5**

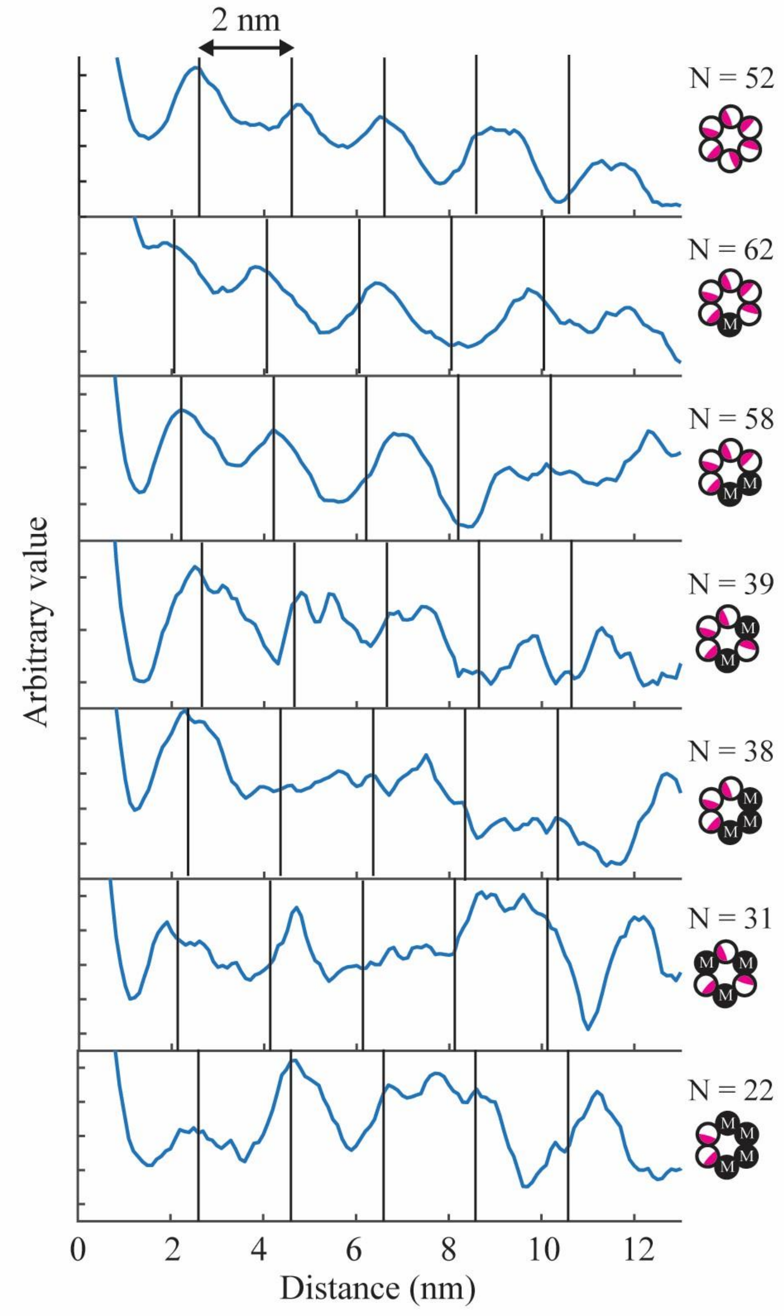


**Supplementary Figure 6**

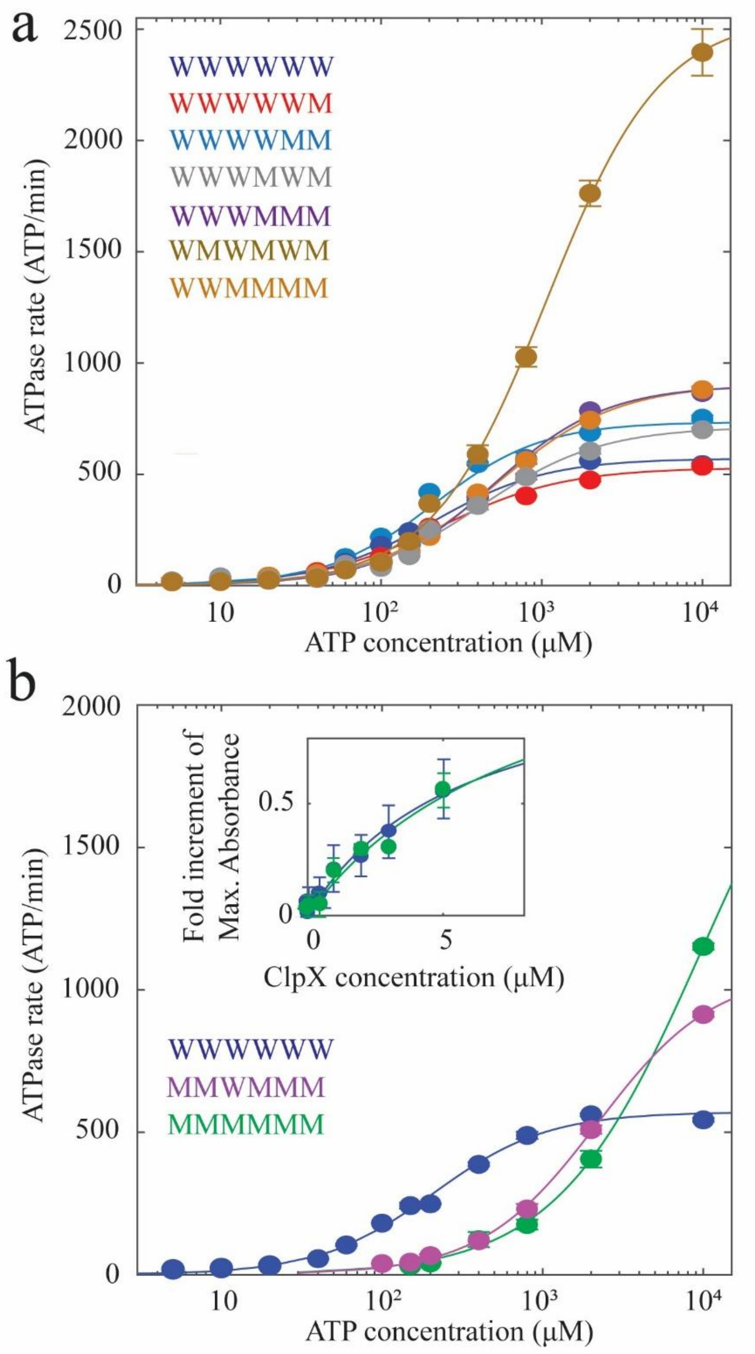


**Supplementary Figure 7**

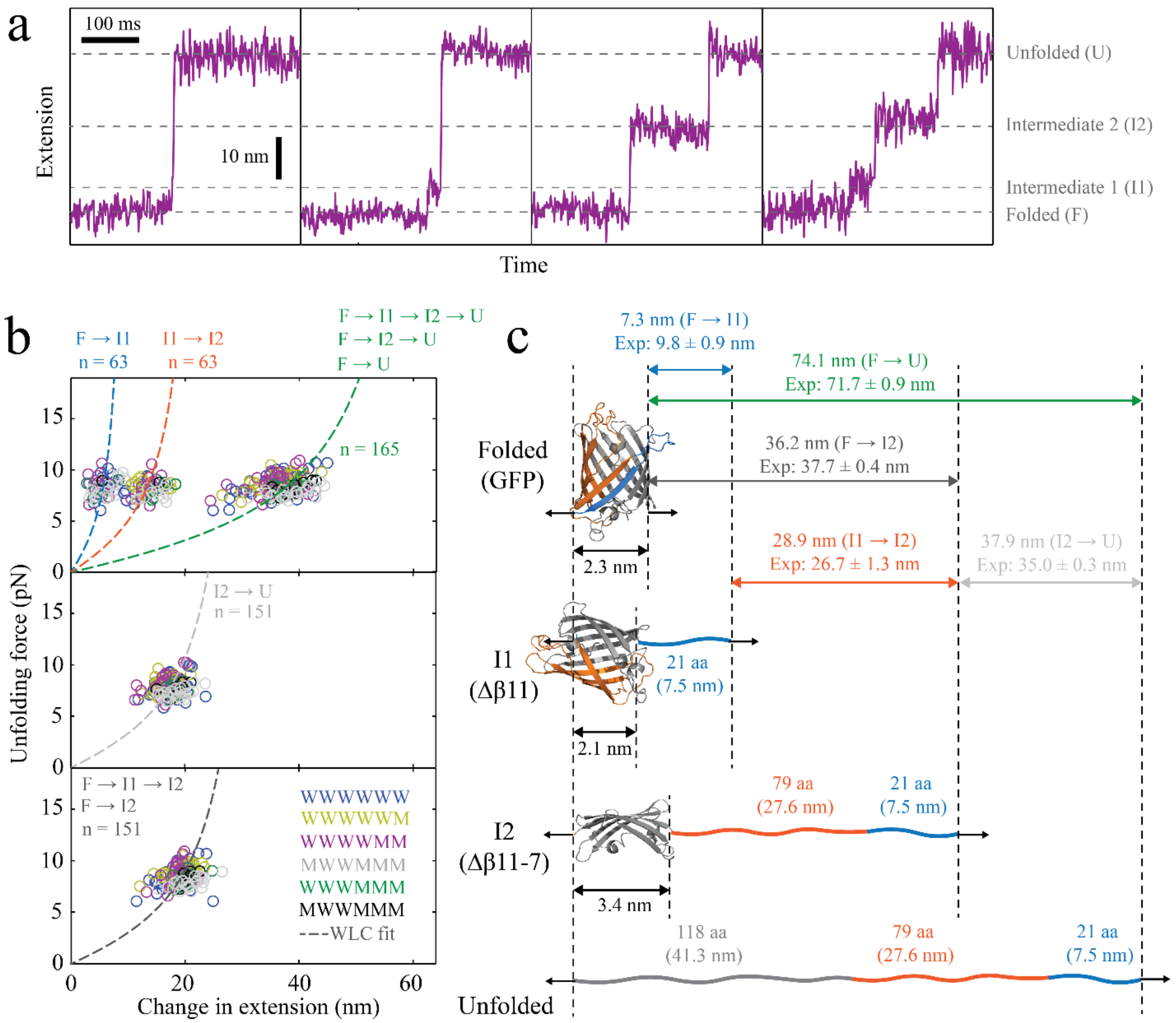


**Supplementary Figure 8**

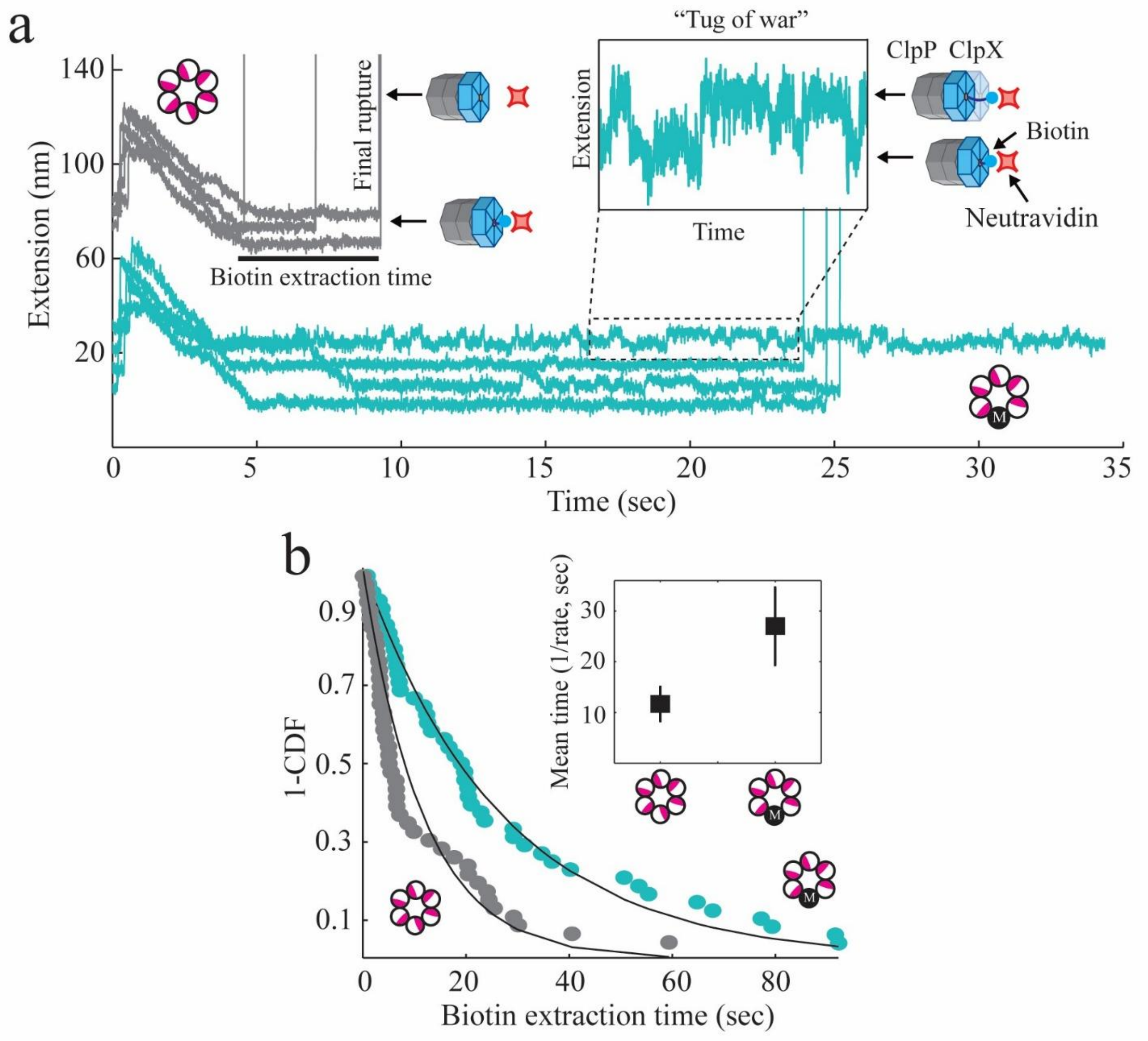


**Supplementary Figure 9**

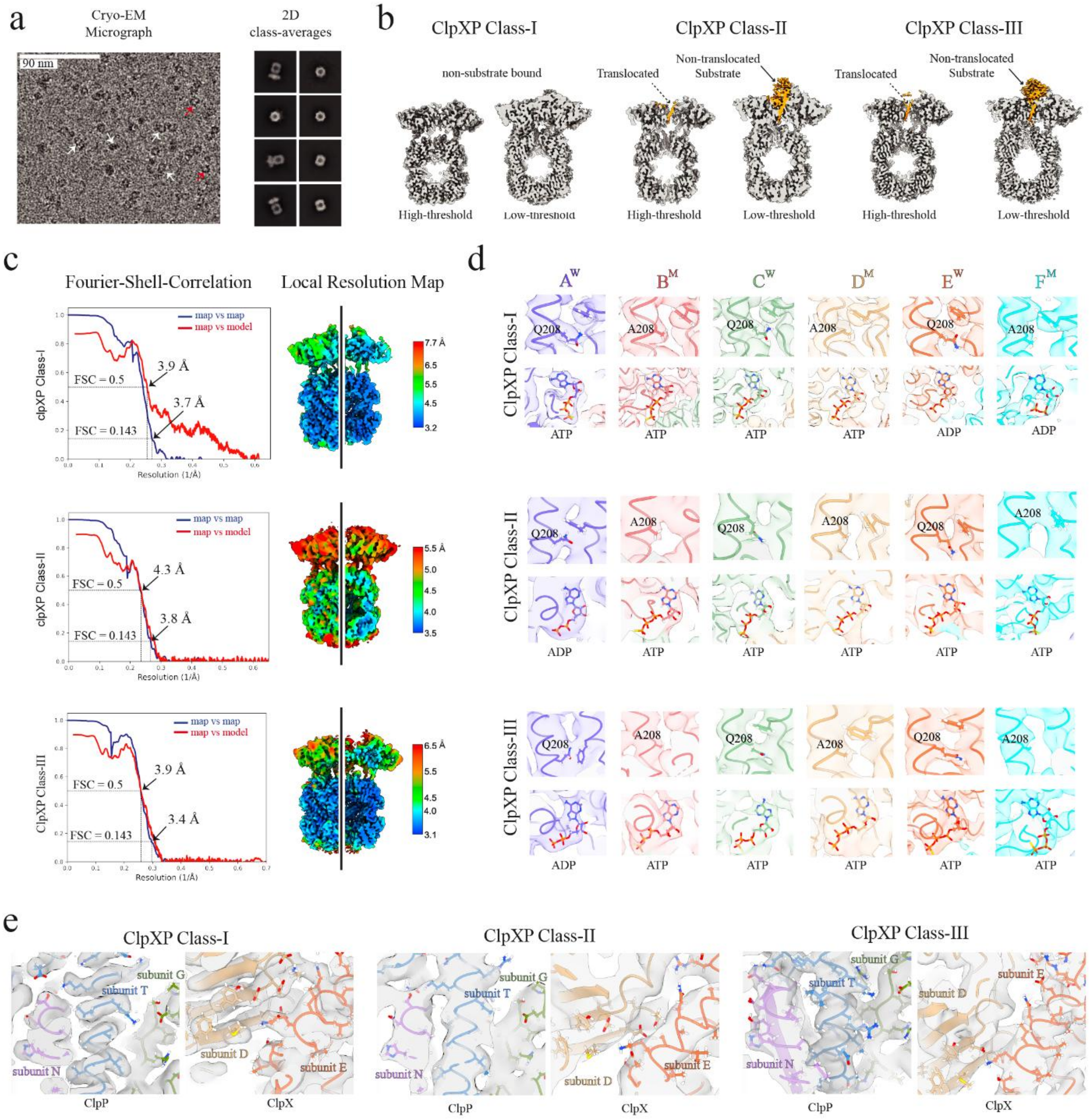


**Supplementary Figure 10**

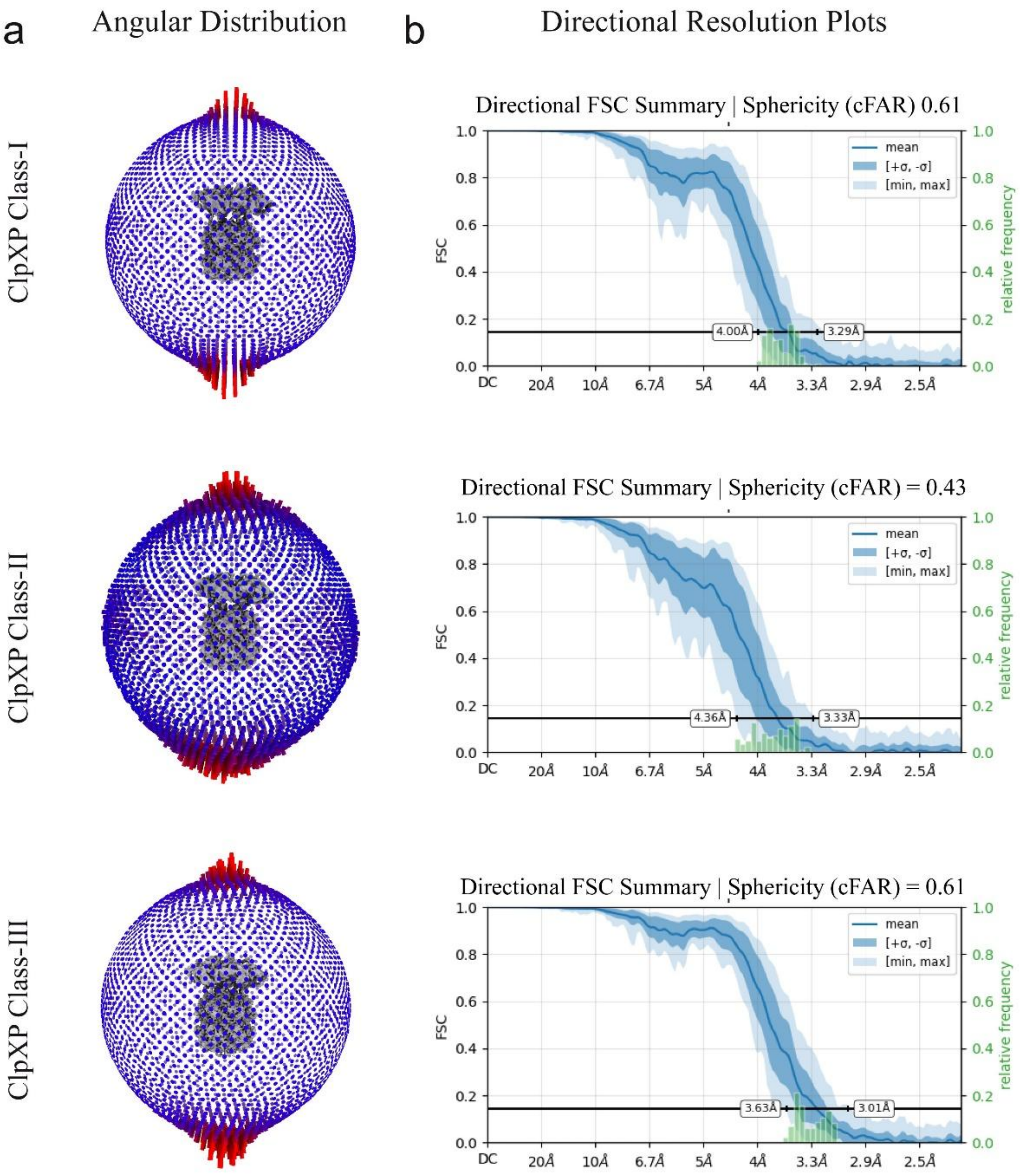


**Supplementary Figure 11**

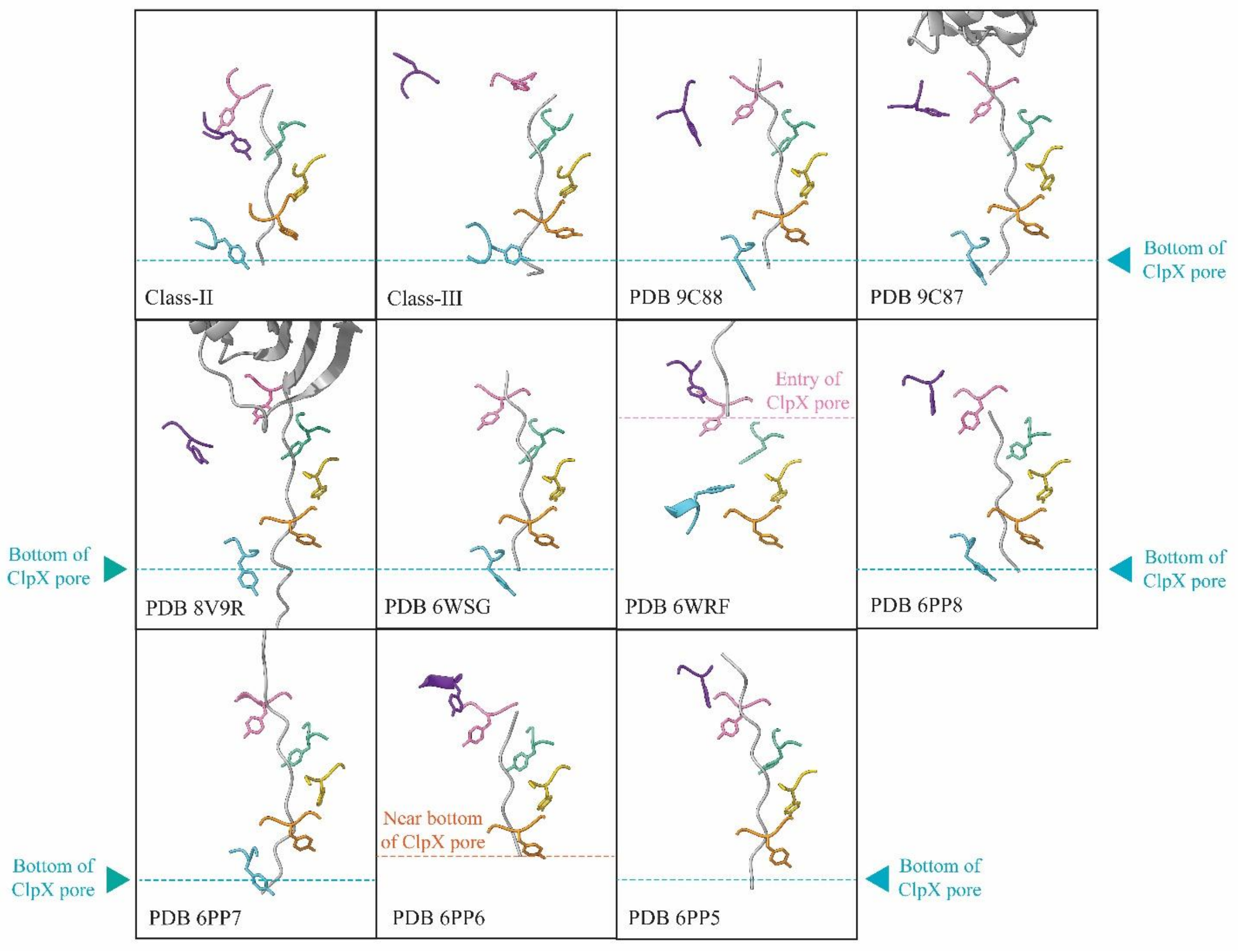


**Supplementary Figure 12**

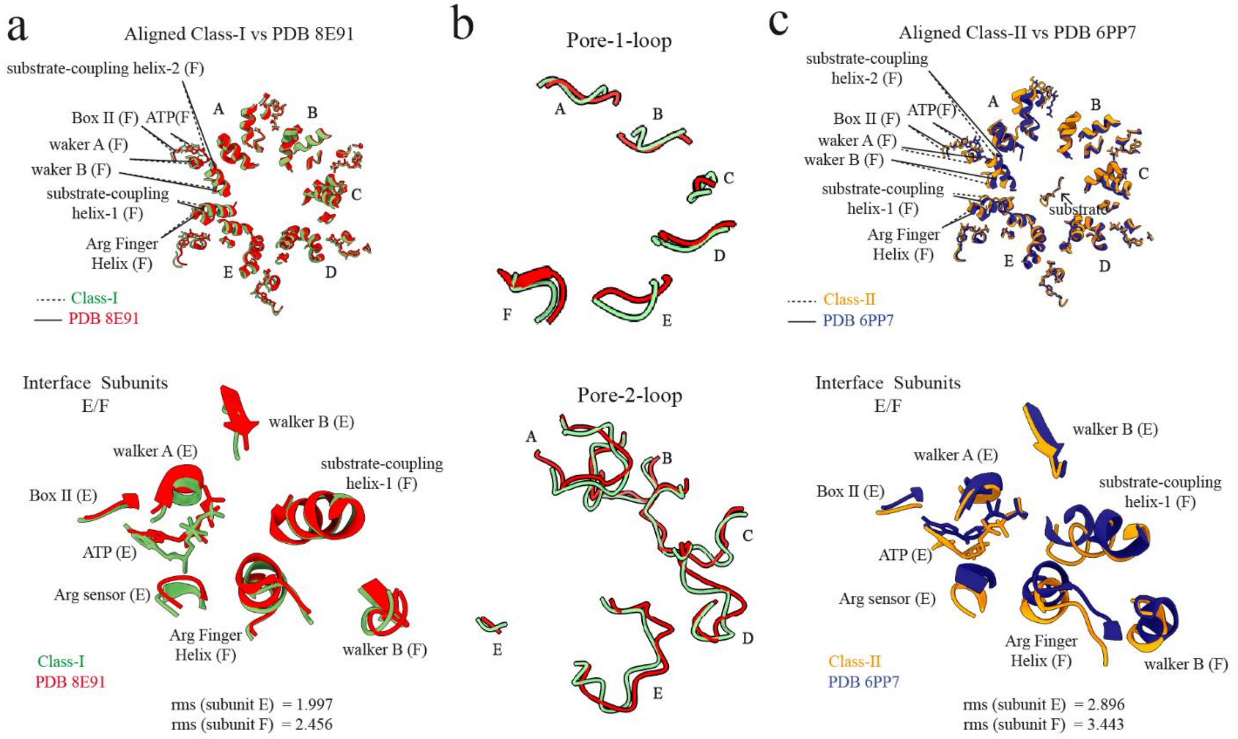


**Supplementary Figure 13**

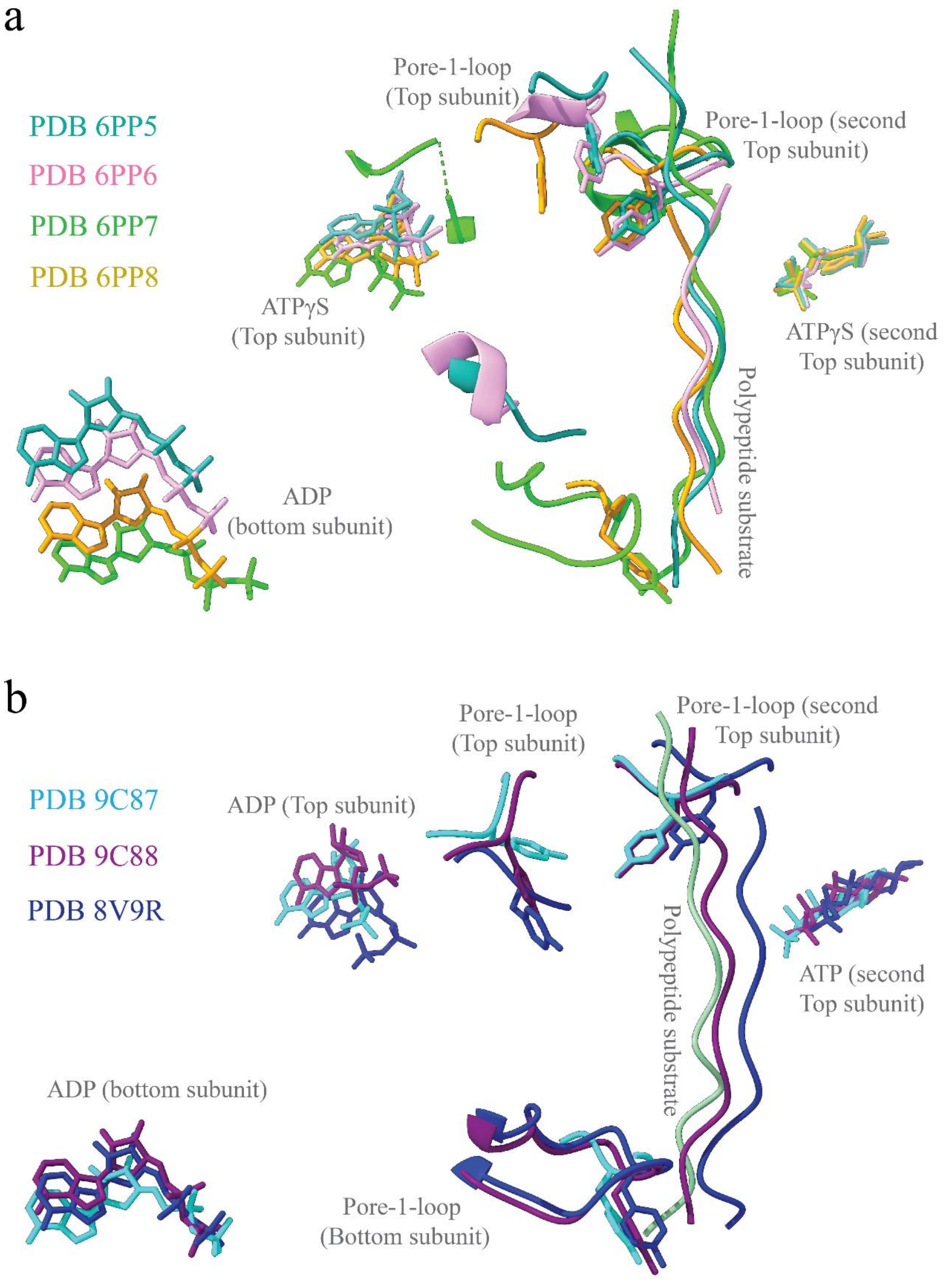


**Supplementary Figure 14**

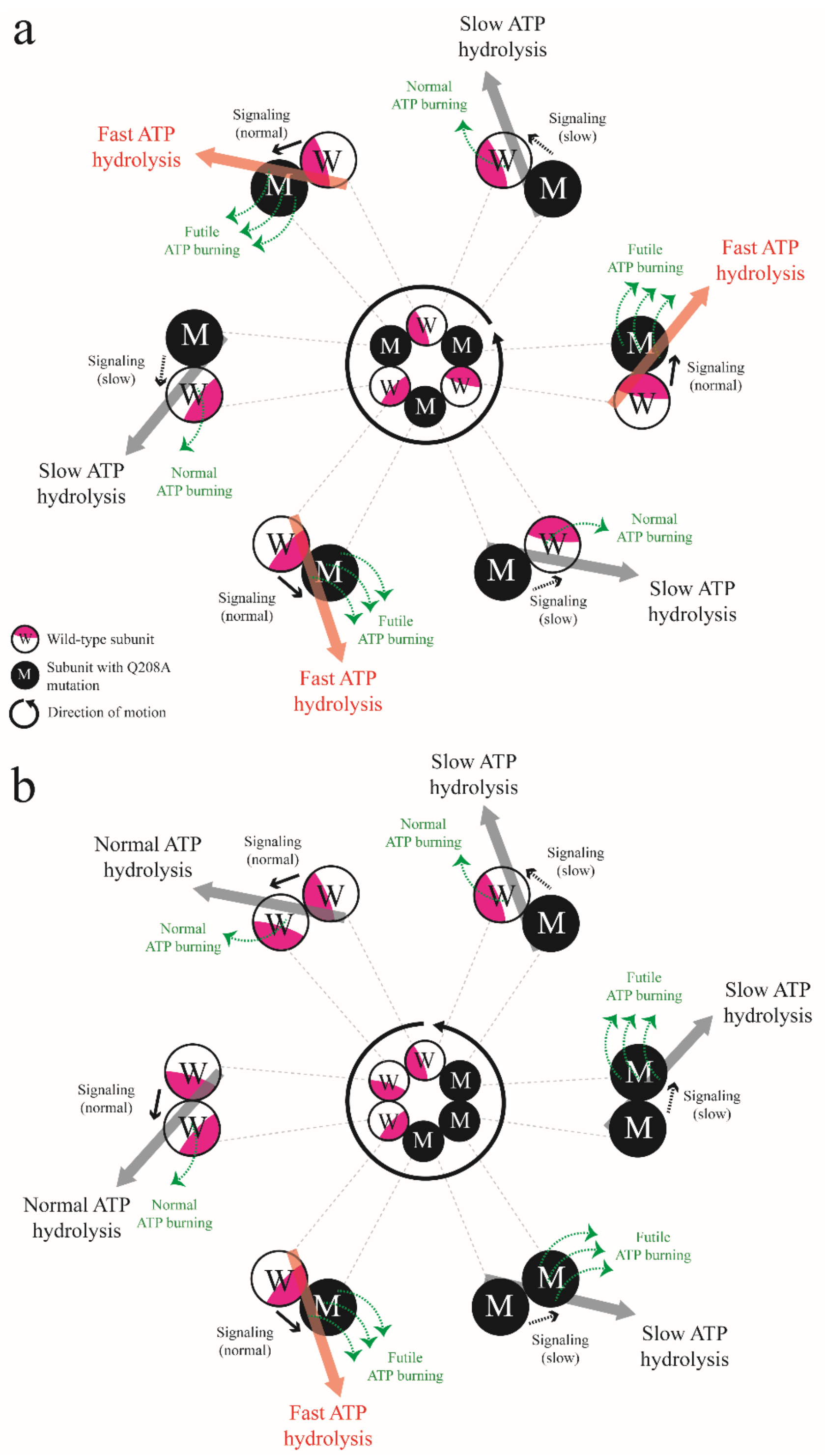


**Supplementary Figure 15**

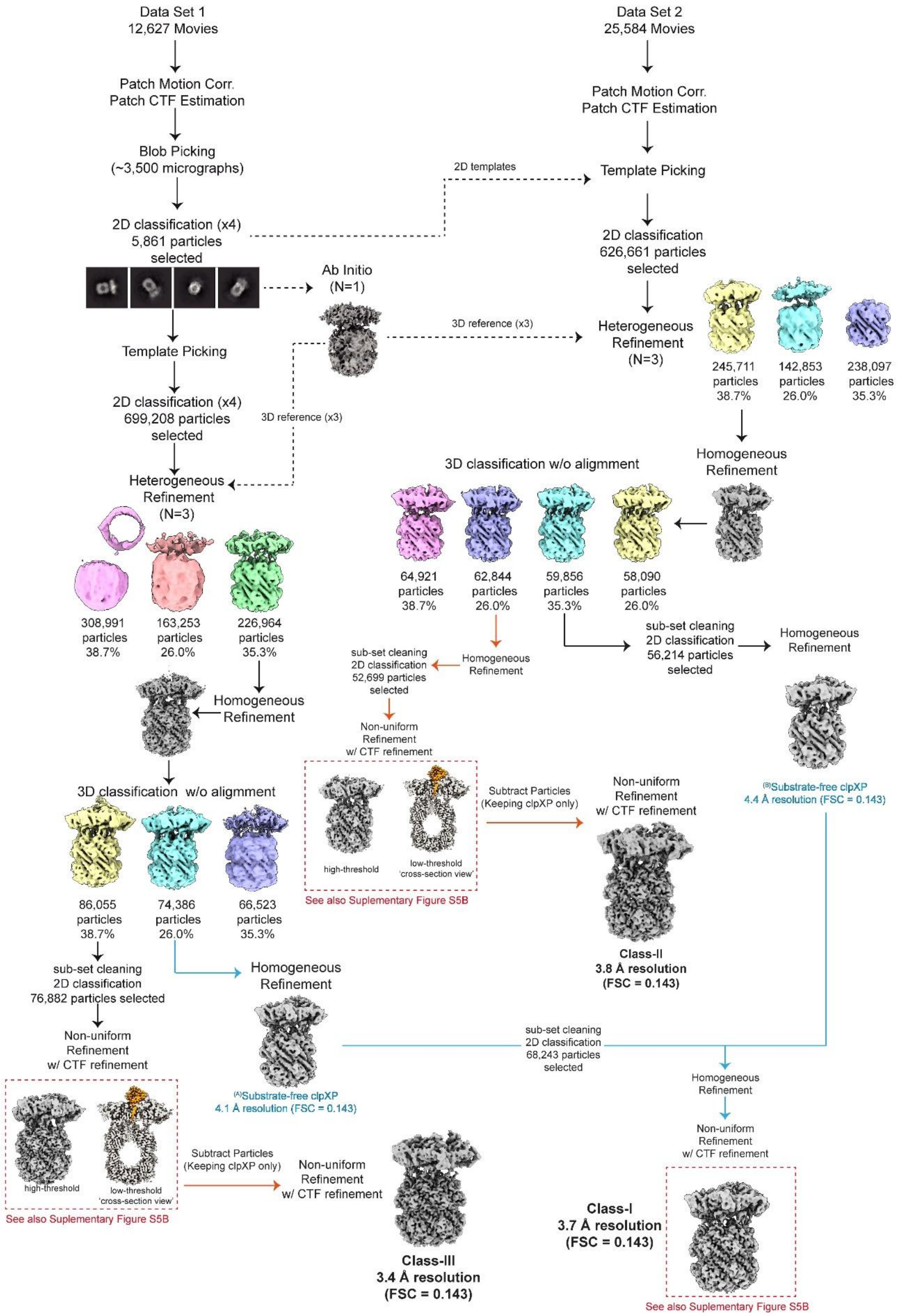


**Supplementary Data 1**

**Supplementary Table 1:** p-values between dwell time distributions using the non-parametric Mann–Whitney U test

| | **WWWWWM** | **WWWWMM** | **WWWMMM** | **WWMMMM** | **WWWMWM** | **WMWMWM** |
|---|---|---|---|---|---|---|
| **WWWWWW** | 1.01E-07 | 9.35E-20 | 1.15E-61 | 1.05E-73 | 6.29E-56 | 2.65E-41 |
| **WWWWWM** | | 7.38E-05 | 7.55E-33 | 1.35E-44 | 1.96E-28 | 6.25E-21 |
| **WWWWMM** | | | 3.58E-16 | 6.78E-26 | 1.07E-12 | 2.28E-09 |
| **WWWMMM** | | | | 0.002534 | 0.307033* | 0.288828* |
| **WWMMMM** | | | | | 4.26E-05 | 0.000157 |
| **WWWMWM** | | | | | | 0.915459* |
| **WMWMWM** | | | | | | |

*Square in orange are the p-values > 0.5, meaning there is no significant difference between those two distributions.

**Supplementary Table 2.** Cryo-EM data collection, refinement and validation Statistics

| | | | |
|---|---|---|---|
| **Data collection** | | | |
| Microscope | Talos Arctica | | |
| Voltage (kV) | 200 | | |
| Detector | Gatan K3 | | |
| Acquisition mode | Super-resolution | | |
| Physical Pixel Size (Å) | 1.141 | | |
| Defocus range (μm) | –0.8 to –2.0 | | |
| Total electron exposure ($e^{-}$ Å$^{-2}$) | 50 | | |
| | ClpXP Class-I (EMD-70686 PDB 9OPA) | ClpXP Class-II (EMD-70772 PDB 9ORD) | ClpXP Class-III (EMD-7086 PDB 9OTX) |
| **Reconstruction** | | | |
| Final particles images (no.) | 124,457 | 52,669 | 76,882 |
| Global Resolution (Å) | 3.7 | 3.8 | 3.4 |
| FSC threshold | 0.143 | 0.143 | 0.143 |
| Map sharpening B-factor (Å$^2$) | 3.2 – 7.7 | 3.5 – 5.5 | 3.1 – 5.5 |
| **Model Composition** | | | |
| Non-hydrogen atoms | 37,256 | 37,230 | 37,043 |
| Protein residues | 4,783 | 4,783 | 4,759 |
| Ligands | | | |
| ATPγS/ATP | 4 | 5 | 5 |
| ADP | 2 | 1 | 1 |
| Mg | 3 | 3 | 3 |
| **Root Mean Square (RMS) Deviations** | | | |
| Bond lengths (Å) | 0.006 | 0.004 | 0.003 |
| Bond angle (°) | 0.732 | 0.677 | 0.672 |
| **B-factors (Å$^2$)** | | | |
| Protein | 75.86 | 175.1 | 125.52 |
| Ligands | 89.43 | 168.02 | 138.2 |
| **Ramachandran Plot** | | | |
| Favored (%) | 98.86 | 98.16 | 98.51 |
| Allowed (%) | 1.14 | 1.6 | 1.38 |
| Outliers (%) | 0 | 0.23 | 0.11 |
| **MolProbity** | | | |
| Clash score | 4.44 | 5.06 | 9.27 |
| Rotamer outliers (%) | 0 | 0.95 | 1.31 |
| Overall score | 1.22 | 1.27 | 1.58 |

**Supplementary Table 3**: Q-scores per subunit for the ClpXP Class-I, -II and -III structures

| Chain | Q-score | | |
|---|---|---|---|
| | **ClpXP class-I** | **ClpXP class-II** | **ClpXP class-III** |
| A | 0.40 | 0.31 | 0.30 |
| B | 0.46 | 0.34 | 0.39 |
| C | 0.45 | 0.37 | 0.44 |
| D | 0.46 | 0.38 | 0.46 |
| E | 0.43 | 0.37 | 0.46 |
| F | 0.35 | 0.32 | 0.37 |
| G | 0.54 | 0.49 | 0.58 |
| H | 0.58 | 0.52 | 0.62 |
| I | 0.59 | 0.52 | 0.64 |
| J | 0.58 | 0.52 | 0.63 |
| K | 0.58 | 0.53 | 0.64 |
| L | 0.59 | 0.52 | 0.64 |
| M | 0.56 | 0.52 | 0.63 |
| N | 0.57 | 0.52 | 0.63 |
| O | 0.53 | 0.50 | 0.58 |
| P | 0.54 | 0.50 | 0.59 |
| Q | 0.54 | 0.50 | 0.58 |
| R | 0.54 | 0.50 | 0.58 |
| S | 0.54 | 0.50 | 0.57 |
| T | 0.55 | 0.49 | 0.58 |

**Supplementary Table 4:** Class-II and Class-III vs previous cryo-EM structures with substrate within the ClpX pore

| Reference | PDB | Nucleotide* | State** | Walker-B mutation/ATPγS | Force affected*** | δ (Å)⁺ |
|---|---|---|---|---|---|---|
| This study (Class-II) | 9ORD | DTTTTT | Pre | No/Yes | Yes | 5.4 |
| This study (Class-III) | 9OTX | DTTTTT | Pre | No/No | Yes | 26.3 |
| Ghanbarpour *et al.*,[2] | 8V9R | DTTTDD | Pre/post | No/No | No | 13.5 |
| Ghanbarpour *et al.*,[2] | 9C87 | DTTTDD | Pre/post | No/No | No | 8.5 |
| Ghanbarpour *et al.*,[2] | 9C88 | DTTTDD | Pre/post | No/No | No | 10.9 |
| Fei *et al.,*[1] | 6PP8 | TTTTTD | Post | Yes/Yes | No | 4.8 |
| Fei *et al.,*[1] | 6PP7 | DTTTTT | Pre | Yes/Yes | No | N.D[++] |
| Fei *et al.,*[1] | 6PP6 | TTTTTD | Post | Yes/Yes | No | N.D[++] |
| Fei *et al.,*[1] | 6PP5 | TTTTTD | Post | Yes/Yes | No | N.D[++] |
| Ripstein *et al.*,[3] | 6VFS | TTTTDD | Post | Yes/No | No | 10.6[+++] |
| Ripstein *et al.*,[3] | 6VFX | TTTTTD | Post | Yes/No | No | 7.6 |

* The nucleotide occupancy of subunits from top to bottom of the ClpX ring spiral. D, denotes ADP, whereas T is ATP.

** The state of the ClpX conformation relative to the power stroke according to the PA/LS model.

*** If the generation of force by ClpX is being affected somewhat and if it was experimentally measured.

[+] The distance between the oxygen of Tyr153 OH-group from the α-carbon of its closest substrate residue located in the ClpX pore.

[++] Not determined because Tyr153 of top subunit (pre-power stroke) was not resolved in PDB 6PP7. Also, Tyr153 of bottom subunit (post-power stroke) was not resolved in PDB 6PP6 and PDB 6PP5.

[+++] This value seems away from the substrate, but the substrate was not solved all the way to the bottom of the ClpX pore.

**Supplementary Table 5:** Model of intersubunit communication and mechanochemical coupling

| | | | | | | | |
|---|---|---|---|---|---|---|---|
| **WMWMWM** | **$k_h \leftarrow k_s$** | Normal ← Slow | Fast ← Normal | Normal ← Slow | Fast ← Normal | Normal ← Slow | Fast ← Normal |
| | **Pair** | W←M | M←W | W←M | M←W | W←M | M←W |
| | **$k_{ATP}$** | Slow | Fast | Slow | Fast | Slow | Fast |
| | **Coupling** | Effective | Futile | Effective | Futile | Effective | Futile |
| **WWWMMM** | **$k_h \leftarrow k_s$** | Normal ← Normal | Normal ← Normal | Normal ← Slow | Fast ← Slow | Fast ← Slow | Fast ← Normal |
| | **Pair** | W←W | W←W | W←M | M←M | M←M | M←W |
| | **$k_{ATP}$** | Normal | Normal | Slow | Slow | Slow | Fast |
| | **Coupling** | Effective | Effective | Effective | Futile | Futile | Futile |
| **WWMMMM** | **$k_h \leftarrow k_s$** | Fast ← Normal | Normal ← Normal | Normal ← Slow | Fast ← Slow | Fast ← Slow | Fast ← Slow |
| | **Pair** | M←W | W←W | W←M | M←M | M←M | M←M |
| | **$k_{ATP}$** | Fast | Normal | Slow | Slow | Slow | Slow |
| | **Coupling** | Futile | Effective | Effective | Futile | Futile | Futile |
| **WWWWWW** | **$k_h \leftarrow k_s$** | Normal ← Normal | Normal ← Normal | Normal ← Normal | Normal ← Normal | Normal ← Normal | Normal ← Normal |
| | **Pair** | W←W | W←W | W←W | W←W | W←W | W←W |
| | **$k_{ATP}$** | Normal | Normal | Normal | Normal | Normal | Normal |
| | **Coupling** | Effective | Effective | Effective | Effective | Effective | Effective |
